# Leveraging Gene Co-expression Patterns to Infer Trait-Relevant Tissues in Genome-wide Association Studies

**DOI:** 10.1101/705129

**Authors:** Lulu Shang, Jennifer A. Smith, Xiang Zhou

## Abstract

Genome-wide association studies (GWASs) have identified many SNPs associated with various common diseases. Understanding the biological functions of these identified SNP associations requires identifying disease/trait relevant tissues or cell types. Here, we develop a network method, CoCoNet, to facilitate the identification of trait-relevant tissues or cell types. Different from existing approaches, CoCoNet incorporates tissue-specific gene co-expression networks constructed from either bulk or single cell RNA sequencing (RNAseq) studies with GWAS data for trait-tissue inference. In particular, CoCoNet relies on a covariance regression network model to express gene-level effect sizes for the given GWAS trait as a function of the tissue-specific co-expression adjacency matrix. With a composite likelihood-based inference algorithm, CoCoNet is scalable to tens of thousands of genes. We validate the performance of CoCoNet through extensive simulations. We apply CoCoNet for an in-depth analysis of four neurological disorders and four autoimmune diseases, where we integrate the corresponding GWASs with bulk RNAseq data from 38 tissues and single cell RNAseq data from 10 cell types. In the real data applications, we show how CoCoNet can help identify specific glial cell types relevant for neurological disorders and identify disease-targeted colon tissues as relevant for autoimmune diseases. Our results also provide empirical evidence supporting one hypothesis of the omnigenic model: that trait-relevant gene co-expression networks underlie disease etiology.

## Introduction

Genome-wide association studies (GWASs) have identified many SNPs associated with common diseases and disease related complex traits. However, over 90% of these identified associations reside in noncoding regions and have unknown biological function [1]. Characterizing the biological functions of these identified associations requires the identification of trait-relevant tissues, as the SNP effects on most traits likely act in a tissue-specific fashion [2, 3]. For example, it is well recognized that brain-specific SNP effects underlie many brain related diseases such as psychiatric disorders [4–7]. For most complex traits, however, their trait-relevant tissues are often obscure. As a result, identifying trait-relevant tissues from GWAS becomes an important first step towards understanding disease etiology and the genetic basis of phenotypic variation [8–15].

Recent development of RNA sequencing (RNAseq) technology, including both bulk RNAseq and scRNAseq, have provided complementary information for the inference of disease relevant tissues. These RNAseq studies produce accurate gene expression measurements both at a genome-wide scale and in a tissue specific or cell type specific fashion. For example, the Gene-Tissue Expression (GTEx) project performs bulk RNAseq to collect gene expression measurements from hundreds of individuals across ~50 tissues [16]. More recently, various scRNAseq studies are being performed to collect cell type specific gene expression measurements on tens of thousands of cells from various tissues and organs [17]. Such tissue specific and cell type specific expression measurements collected from bulk RNAseq and scRNAseq provide valuable information for inferring disease relevant tissue types. Indeed, methods have been developed to identify genes that are specifically expressed in a particular tissue or cell type to construct tissue specific or cell type specific annotations at the gene level, which are further integrated into GWASs to infer disease-relevant tissues or cell types [18, 19]. However, these previous methods have ignored an important biological feature of gene expression data; that is, genes are interconnected with each other and are co-regulated together. Such gene co-expression patterns occur in a tissue specific or cell type specific fashion [8]. Certain gene co-expression sub-networks have been shown to contain valuable information for predicting gene-level association effect sizes on diseases in GWASs [8, 20–22]. In addition, genes with high network connectivity are enriched for heritability of common diseases and disease related traits [23]. Indeed, one key hypothesis in the recent omnigenic model states that tissue-specific gene networks underlie the etiology of various common diseases [24]. Therefore, it is important to develop statistical methods that can take advantage of tissue-specific topological connections contained in tissue specific gene co-expression networks to facilitate the inference of disease tissue relevance.

Here, we make such a first attempt to integrate GWAS data with gene co-expression patterns obtained from gene expression studies, through developing a statistical method for the inference of trait-relevant tissues. To do so, for a given trait, we treat the gene-level association statistics obtained from GWAS as the outcome variable and treat the tissue-specific adjacency matrices inferred from gene expression studies as input matrices. We examine one tissue at a time and model the gene-level association statistics as a function of the tissue-specific adjacency matrix. Afterwards, we identify trait-relevant tissues through likelihood-based inference. To accompany our model, we also develop a composite likelihood-based algorithm, which is computationally efficient and ensures result robustness in the presence of substantial noise in the estimated tissue-specific gene adjacency matrices. We refer to our method as *Co*mposite likelihood-based *Co*variance regression *Net*work model, or CoCoNet. We demonstrate the effectiveness of our method through simulations. We apply our method for an in-depth analysis of four autoimmune diseases and four neurological disorders, through integrating the corresponding GWASs with bulk RNAseq data from 38 tissues and single cell RNAseq data from 10 cell types. In the real data applications, we show how our method and analysis can help reveal new biological insights and help provide empirical evidence supporting an important hypothesis from the omnigenic model, namely that trait-relevant gene co-expression networks underlie disease etiology.

## Materials and Methods

### Covariance Regression Network Model

We aim to leverage tissue-specific gene co-expression networks to infer trait relevant tissues through integrating GWAS and gene expression studies. To do so, we perform gene-centric analysis and focus on a common set of *m* genes that are measured in both GWAS and gene expression studies. For these genes, we obtain an *m*-vector of summary statistics in terms of gene-level effect sizes from GWAS for the trait of interest and denote the vector as ***y*** = (*y*_1_, …, *y_m_*)^T^. We also obtain tissue-specific gene expression measurements from gene expression studies on multiple tissues. For each tissue in turn, we construct an *m* by *m* gene-gene adjacency matrix to represent the co-expression network. Such adjacency matrix is constructed based on gene expression measurements, paired with prior gene-gene interaction information obtained from external data sources (more details in the following subsections). We denote the constructed tissue-specific symmetric adjacency matrix as ***A*** = (*a_ij_*), where its *ij*’th element *a_ij_*. is one if gene *i* is connected to gene *j* in the network and is zero otherwise. We set *a_ij_* to be zero for any 1 ≤ *i* ≤ *m* to ensure the absence of self-loops [25]. In addition, we denote 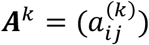 as the *k*-th power of ***A***, for any integer *k*. It can be easily shown that 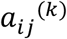 is the number of *k*-paths linking from gene *i* to gene *j* in the co-expression network, where *k*-paths are any paths of length *k*. For example, when 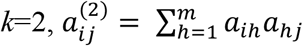, where *a_ih_a_hj_*. is one only when there is a link connecting the three genes *i* – *h* – *j* and is zero otherwise. We also set 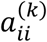 to be zero for *k* ≥ 1. When *k* = 0, we define *A^0^* = *I*, which is an m-dimensional identity matrix.

We reason that, in the trait-relevant tissue, if two genes share similar functionality, then these two genes will likely have similar effects on the trait of interest. In contrast, in the trait irrelevant tissue, two genes sharing similar functionality would not be strongly predictive of their effect size similarity on the trait of interest. Therefore, the prediction ability of the adjacency matrix ***A*** on the gene-level effect size *y_i_* would be an effective indication on whether the examined tissue is relevant to trait or not. To capture such intuition, we use the Covariance Regression Network Model [26] to model the relationship between ***A*** and ***y***. Specifically, we consider

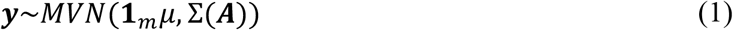

where ***1**_m_* is an m-vector of 1s; *μ* is the intercept; MVN denotes multivariate normal distribution; and Σ(***A***) is the covariance of ***y*** that is a function of the adjacency matrix ***A***. Following [26], we consider the use of polynomial matrix functions for the construction of Σ(***A***) by setting 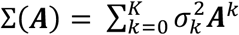, where *K* is the maximum number of paths linking between two genes considered in the model and is treated as a pre-fixed parameter. Intuitively, 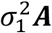 captures the gene-level effect size correlation due to direct connections among genes, while 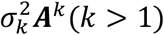 captures the gene-level effect size correlation due to indirect connections among genes (i.e. gene-gene connection through other genes).

The above model can be fitted through a standard maximum likelihood inference procedure. However, parameter estimation through the standard maximum likelihood inference procedure is computationally inefficient as it scales cubically with the number of genes *m*. To enable scalable computation, we consider the composite likelihood approach for inference [27]. Specifically, instead of working on the joint likelihood specified in equation (1), we consider pairs of genes one at a time. For each pair of genes *i* and *j*, we consider the pair-wise likelihood 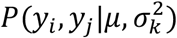 as

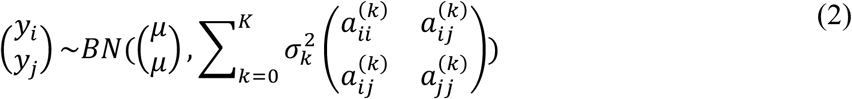

where *BN* denotes a bivariate normal distribution. With the model specified in equation (2), we can obtain the corresponding log composite likelihood as:

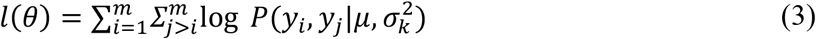

We use the Nelder-Mead method implemented in the optim function in R to obtain the composite likelihood estimates 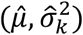 that maximizes the above composite likelihood. Our inference algorithm scales only quadratically with the number of genes, and, with a small *K*, can analyze each trait-tissue pair with 10,000 genes in a few minutes.

Besides scalable computation, we note that the composite likelihood-based inference algorithm also ensures result robustness with respect to model mis-specifications. Specifically, instead of making the strong assumption that the *m*-vector of ***y*** jointly follows a multivariate Gaussian distribution, our composite likelihood only needs to make a much weaker assumption that each pair (*y_i_, y_j_*) follows a bivariate normal distribution. As a result, the composite likelihood-based algorithm can be robust to various model misspecifications, relieving a potential concern in real data applications, where the tissue specific adjacency matrices may be estimated with a substantial estimation noise.

Finally, in the above model, the number of covariance matrices used, *K*, is treated as a fixed parameter. In the real data applications, we explored a few different choices of *K* in the range between one and four. We found that models with small *K* values are often preferred based on Bayesian Information Criterion (BIC): in all of the trait-tissue pairs, a model with *K* = 1 has the smallest BIC (S1 Fig). The lower BIC in models with small *K* is presumably because the direct connections described in ***A*** contain most information for predicting the correlation among gene-level effect sizes. Therefore, due to convenience and practical effectiveness, we focus on modeling with *K* = 1 in all our simulations and real data applications. However, we note that our CoCoNet software implementation is general: it applies to any pre-specified *K* and can also perform model selection to determine the optimal choice of K. When *K* = 1, we define 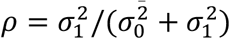, which represents the relative signal strength of gene co-expression pattern on gene-level effect sizes ***y***.

We refer to the above model as *Co*mposite likelihood-based *Co*variance regression *Net*work model, or CoCoNet. With the composite likelihood inference algorithm, we examine one tissue at a time and obtain the maximum composite likelihood. Afterwards, we rank the tissues based on the maximum composite likelihood and select the tissue with the highest likelihood as the most trait-relevant tissue. Because of the composite likelihood approach, CoCoNet is computationally efficient and can analyze each trait-tissue pair in real data in minutes (S1 Table). The CoCoNet method is implemented as an R package, which, together with all processed data and scripts to reproduce the results in the paper, are freely available at www.xzlab.org/software.html.

### Simulations

We performed simulations to examine the effectiveness of our method. To do so, we first randomly selected 10 tissues and 1,000 genes from the genotype-tissue expression (GTEx) study. We then used these gene expression data to construct tissue-specific gene adjacency matrices, with which we further simulated gene level effects sizes as outcomes. Specifically, for each tissue in turn, we first categorized the selected 1,000 genes into non-overlapping gene clusters using the *k*-means clustering algorithm. The number of clusters for each tissue was determined based on BIC and ranged from 10 to 15 across tissues. Based on the gene clusters, we constructed the tissue-specific gene adjacency matrix as a block diagonal matrix: two genes are adjacent to each other if both belong to the same inferred cluster and are not adjacent to each other otherwise. Note that we constructed the gene adjacency matrices in a simpler way in the simulations than in the real data (details of how gene adjacency matrices are constructed in the real data are provided in the following section), to ensure that the covariance matrices are positive definite in the simulations so that we can easily simulate outcome variables from multivariate normal distributions. Next, we denoted one of the 10 tissues as the trait-relevant tissue and used its tissue-specific adjacency matrix to simulate our outcome variables. Afterwards, we fit data using the 10 tissues one at a time to identify the trait-relevant tissue. We examined three main simulation scenarios, each consisting of multiple parameter settings. We performed 100 simulation replicates for each parameter setting. We computed power as the percent of replicates where the true trait relevant tissue is correctly identified.

#### Scenario I

The first simulation scenario is based on our model and assumes that we can directly observe the gene adjacency matrices for all tissues. Here, in each simulation replicate, we randomly designated one tissue out of the 10 tissues to be the trait-relevant tissue. We simulated the outcome variables through a multivariate normal distribution with mean zero and covariance matrix as 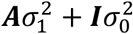, where ***A*** is the adjacency matrix from the designated trait-relevant tissue. In the simulations, we set 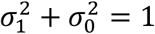 and varied 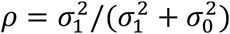 in the range of 0 to 0.09 to examine the influence of signal strength on power. Note that *ρ* = 0.02 is close to the median estimate in the real data applications. In each replicate, we applied our model to examine one tissue at a time, treated the tissue-specific adjacency matrices as observed, and selected among the 10 tissues the one with the highest log likelihood (equivalently the lowest BIC) as the trait-relevant tissue.

#### Scenario II

The simulation scenario II is similar to scenario I, except that we were unable to observe the true adjacency matrices. Instead, we were able to observe only a noisy version of the adjacency matrices. Specifically, we simulated the data using the true tissue-specific adjacency matrix as done in scenario I. However, when we fitted data, we were only provided with the observed tissue-specific adjacency matrices that were a noisy version of the truth. To generate these observed adjacency matrices, for each tissue in turn, we randomly converted a proportion *p* of the adjacent gene pairs in the true adjacency matrix to be nonadjacent, and randomly converted a proportion *q* of nonadjacent gene pairs in the true adjacency matrix to be adjacent. For each value of *p*, we chose the value of *q* so that the total number of adjacent gene pairs was the same between the true adjacency matrix and the observed adjacency matrix. Here, we fixed *ρ* = 0.02, a value close to the median estimate in the real data application; and we varied *p* from 0 to 0.9 to capture an increasingly large measure of noise.

#### Scenario III

The simulation scenario III is similar to scenario II, except that the number of adjacent gene pairs in the observed adjacency matrices is much larger than the number of adjacent gene pairs in the true adjacency matrices, potentially reflecting an over-estimation of the number of adjacent gene pairs that may be observed in some real data applications. The observed adjacency matrices in this scenario was created by adding new adjacent gene pairs to the true adjacency matrices. Specifically, for each tissue in turn, we randomly converted a proportion *q* of nonadjacent gene pairs in the true adjacency matrix to be adjacent in the observed adjacency matrix. The adjacent genes in the true adjacency matrix serve as core genes that truly influence the gene-level association evidence with the complex trait. We again simulated the data using the true adjacency matrix and fitted data using the observed adjacency matrices. Here, we fixed *ρ* = 0.02, a value close to the median estimate in the real application; and we varied *q* from 0 to 0.9 to capture a range of core gene set proportions.

### Real Data Sets

#### Gene-level effect sizes from GWASs

We examined a total of eight disease traits from existing large-scale GWASs. These traits include four neurological disorders and four autoimmune disorders, with GWAS sample sizes ranging from 13,239 to 70,100. The examined neurological disorders include schizophrenia (SCZ) [28], bipolar disorder/schizophrenia (BIPSCZ) [29], bipolar disorder (BIP) [29], and Alzheimer’s disease [30], where BIPSCZ includes combined individuals from SCZ and BIP. The examined autoimmune disorders include primary biliary cirrhosis (PBC) [31], ulcerative colitis (UC) [32], inflammatory bowel disease (IBD) [32], and Crohn’s disease (CD) [32]. We selected these traits because the trait-relevant tissues for both neurological disorders and autoimmune disorders are relatively clear and because these traits are measured on GWASs with at least 12,000 samples, ensuring sufficient power [33]. The information for the summary statistics of eight GWAS traits can be found in S2 Table.

For each of these traits, we first obtained SNP-level summary statistics in the form of marginal z-scores. Following [34], we removed SNPs within the major histocompatibility complex (MHC) region (Chr6: 25Mb-34Mb). We intersected the remaining SNPs from all eight studies to retain a common set of 622,026 SNPs for analysis. Besides SNP-level marginal z-scores, we also obtained individual-level genotypes of 503 European individuals from the 1,000 Genomes Project to serve as a reference panel for linkage disequilibrium (LD) computation [35]. In addition, we obtained location information for 51,014 genes from the GENCODE project [36], and extracted cis-SNPs that reside within 1Mb before the transcription start site (TSS) and within 1Mb after the transcription end site (TES). We focused on 49,015 genes that have at least 10 SNPs, with an average of 438.8 SNPs in each gene (median = 448; min = 10; max = 2,425). Afterwards, with SNP-level statistics and LD information from the reference panel, we obtained gene-level heritability estimates using MQS [37]. We scaled the gene-level heritability estimates by the number of SNPs in each gene. Afterward, we further standardized the scaled values across genes to have a mean of zero and standard deviation of one. These final values are served as the gene level effect sizes for the GWAS trait.

#### Tissue specific gene co-expression networks from GTEx

We obtained tissue-specific gene co-expression networks inferred based on bulk RNAseq data collected on 38 tissues in the GTEx project [38] from https://zenodo.org/record/838734#.XALkry3MxTZ. Details on how these tissue-specific gene coexpression networks were constructed are described in [39]. Briefly, these networks were constructed using the software PANDA (Passing Attributes between Networks for Data Assimilation) [40], relying on information extracted from both gene expression measurements and an existing database that contains both known transcription factor (TF)-target gene interactions and known protein-protein interactions. We intersected the set of genes in the aforementioned database with the set of genes from GWASs to obtain an overlapping set of 25,991 genes for analysis. In the networks constructed through PANDA, the genes are represented as nodes while the connected gene pairs are represented as edges. Each gene and each edge have a specificity score calculated in each tissue. Following [39], we focused our analysis on the edges that are identified to be specific in at least one tissue, and genes that are specific (i.e. with a non-zero value) in at least two tissues (details in [39]). In addition, we included genes that are TF factors based on the database and have at least one connection to genes that we already retained. We focused our analysis on a final set of 5,359 genes. The edge value between a pair of genes represents evidence on whether the given gene pair is connected in the network or not: a strongly positive edge value indicates greater evidence on the existence of gene-gene connection. The edge values across all gene pairs have an approximate mean value of zero and a standard deviation of one. For edge values, we performed a hard thresholding procedure to convert the continuous edge values to binary values. In particular, edges that had a positive edge value and that were specific to at least one tissue type were converted to one; otherwise, they were converted to zero. This way, tissue-specific gene co-expression networks were converted to tissue-specific adjacency matrices used in our model.

#### Cell type specific gene co-expression networks from GTEx

We obtained single cell nucleus sequencing data with droplet technology (DroNc-seq) generated on archived frozen adult human post-mortem tissues from the GTEx project [41]. The GTEx DroNc-seq data contains gene expression measurements on 32,111 genes and 14,963 single cells from adult frozen human hippocampus (Hip, 4 samples) and prefrontal cortex (PFC, 3 samples) from five donors. We retained 16,930 genes that overlapped with genes from the GWASs. For each gene in turn, we transformed the data in the unit of reads per kilobase million (RPKM), and performed log10 transformation on the RPKM values after adding a constant of one. Following [42], we normalized the expression values across genes in each cell to a standard normal distribution, and further normalized the expression values of each gene across cells to a standard normal distribution. All cells in the data were already clustered into 10 cell types in the original paper using the *k*-nearest neighbor (*k*-NN) method [41]. These 10 cell types are exPFC, which consists of excitatory glutamatergic neurons in the prefrontal cortex; GABA, which consists of GABAergic interneurons; exCA, which consists of excitatory pyramidal neurons in the hippocampal Cornu Ammonis (CA) region; exDG, which consists of excitatory granule neurons from the hippocampal dentate gyrus region; ASC, which consists of astrocytes; MG, which consists of microglia cells; ODC, which consists of oligodendrocytes; OPC, which consists of oligodendrocyte precursor cells; NSC, which consists of neuronal stem cells; and END, which consists of endothelial cells. Following [39], we used the software PANDA to infer cell type specific gene co-expression networks for these 10 different cell types in each of the two donors separately. Because the gene co-expression networks for the same cell type in the two donors are similar to each other, we merged the inferred gene regulatory networks for the same cell type from the two donors together by taking the unions of the two corresponding gene co-expression networks. In the constructed networks, we calculated gene specificity scores (details in [39]) and retained half of the genes with a total specificity score across tissues above the median value. In addition, we retained genes that are either TF factors or have at least one connection to genes we already retained. This way, we retained a total of 8,269 genes for final analysis. We created cell type specific gene co-expression adjacency matrices for these genes based on the inferred coexpression networks following the same procedure described in the previous section.

#### Relevance between traits and tissues/cell types from PubMed search

Following [8], we partially validated the identified trait-relevant tissue/cell types for the GWAS diseases by searching PubMed. We reasoned that, if the tissue or cell type is relevant to the disease of interest, then there would be previous publications studying the disease in the particular tissue or cell type. Therefore, by counting the number of previous publications using the key word pairs of trait and tissue or the key word pairs of trait and cell type, we would have a reasonable quantitative estimate on the relevance between the trait and the corresponding tissue/cell type. To do so, we counted the number of references on the disease and tissue pairs for the 8 GWAS diseases and 38 tissues. In addition, we also counted the number of references on the disease and cell type pairs for the 4 neurological disorders and 10 brain cell types. We used an R package RISmed (https://cran.r-project.org/web/packages/RISmed/index.html) to efficiently count the number of publications in PubMed that contain the names of both the trait and the tissue/cell type either in the abstract or in the title [43]. The keywords of traits and tissues/cell types that we used in searching are listed in the supplementary S3-S5 Tables. For example, for the trait-tissue pair of schizophrenia and cerebellum, we conducted the search by using “Schizophrenia[Title/Abstract] AND cerebellum[Title/Abstract]”, which yielded 946 hits. After obtaining the number of papers on each trait-tissue/cell type pair, for each trait at a time, we further normalized the count data across tissues or across tissues/cell types by calculating the percentage of publications on each tissue or on each tissue/cell type.

#### Inferring trait relevant tissues/cell types with RolyPoly and LDSC

We compared our results with two existing methods that use gene-level annotations to infer trait relevant tissue/cell types: RolyPoly and LD score regression (LDSC). RolyPoly requires input that include GWAS summary statistics, gene expression profiles, an expression data annotation file, and linkage disequilibrium (LD) information. For GWAS summary statistics and gene expression profiles, we used the same input data for RolyPoly as used for our analysis. To make the gene annotation file, we defined a block as a 10kb window centered around each gene’s transcription start site (TSS) as recommended by RolyPoly. For LD information, we used the LD information provided by RolyPoly website, which were based on the 1,000 genomes project phase 3 data and which set SNP pair covariance to be zero if below 0.2. For LDSC, following [44], we calculated t-statistics for each gene being differentially expressed in a given tissue or cell type versus all other tissue/cell types that are not in the same tissue category. For example, for cerebellum, we compared expression in cerebellum samples versus expression in all other samples but excluding the other brain regions. We then selected the top 2,000 tissue/cell type specific genes ranked by t-statistics. For these 2,000 genes, following [44], we annotated SNPs within 100-kb of their transcribed regions to have an annotation value of one and annotated the remaining SNPs to have an annotation value of zero. We then ran LDSC to estimate the heritability enrichment using the SNP annotation for each of the eight GWAS traits separately. Both RolyPoly and LDSC output *p* value for each examined trait-tissue pair, with which we ranked tissues for each trait.

## Results

### Method and Simulations

Our method is described in the Materials and Methods, with technical details provided in the Supplementary Text. Briefly, our method requires both gene-level effect sizes obtained from GWAS on the trait of interest and tissue-specific gene co-expression adjacency matrices inferred from gene expression studies. For a given trait of interest, we examine one tissue at a time and model the gene-level effect sizes for the trait as a function of the gene co-expression matrix using a covariance regression network model. To ensure model robustness and scalable computation, we infer parameters in the covariance regression network model through composite likelihood. We calculate the maximum composite likelihood for each tissue and eventually rank tissues by the corresponding log likelihoods. We refer to our method as CoCoNet, freely available as an R software package.

We first examined the performance of our method through simulations (details in Materials and Methods). Briefly, we created tissue-specific gene co-expression adjacency matrices for 1,000 genes from 10 tissues in GTEx. We randomly selected one tissue as the true trait-relevant tissue and used its gene co-expression adjacency matrix to simulate the gene-level effect sizes for the trait. Afterwards, we examined one trait at a time using CoCoNet and calculated the power to detect the true trait-relevant tissue. We examined a total of three scenarios.

In scenario I, we fit the data using the tissue-specific gene co-expression adjacency matrices. We varied the signal strength parameter *ρ* within the model to examine the estimation accuracy and power of our method across a range of signal strength (Fig 2A). We found that our composite likelihood-based method can estimate model parameters relatively well (S2 Fig A and D, S3 Fig A and D, S4 Fig A and D). For example, when *ρ* = 0.02, which is close to the median estimate in the real data applications, the *ρ* estimates across 100 simulation replicates are centered around 0. 017, with a standard deviation of 0.013. In addition, our method has descent power in detecting the true trait-relevant tissue across a range of *ρ* (Fig 2D). For example, when *ρ* = 0.02, the power of our method is 0.65. The power of our method also increases with increasing *ρ*. For example, the power of our method increases to 0.89 when *ρ* = 0.03.

**Fig 1.**
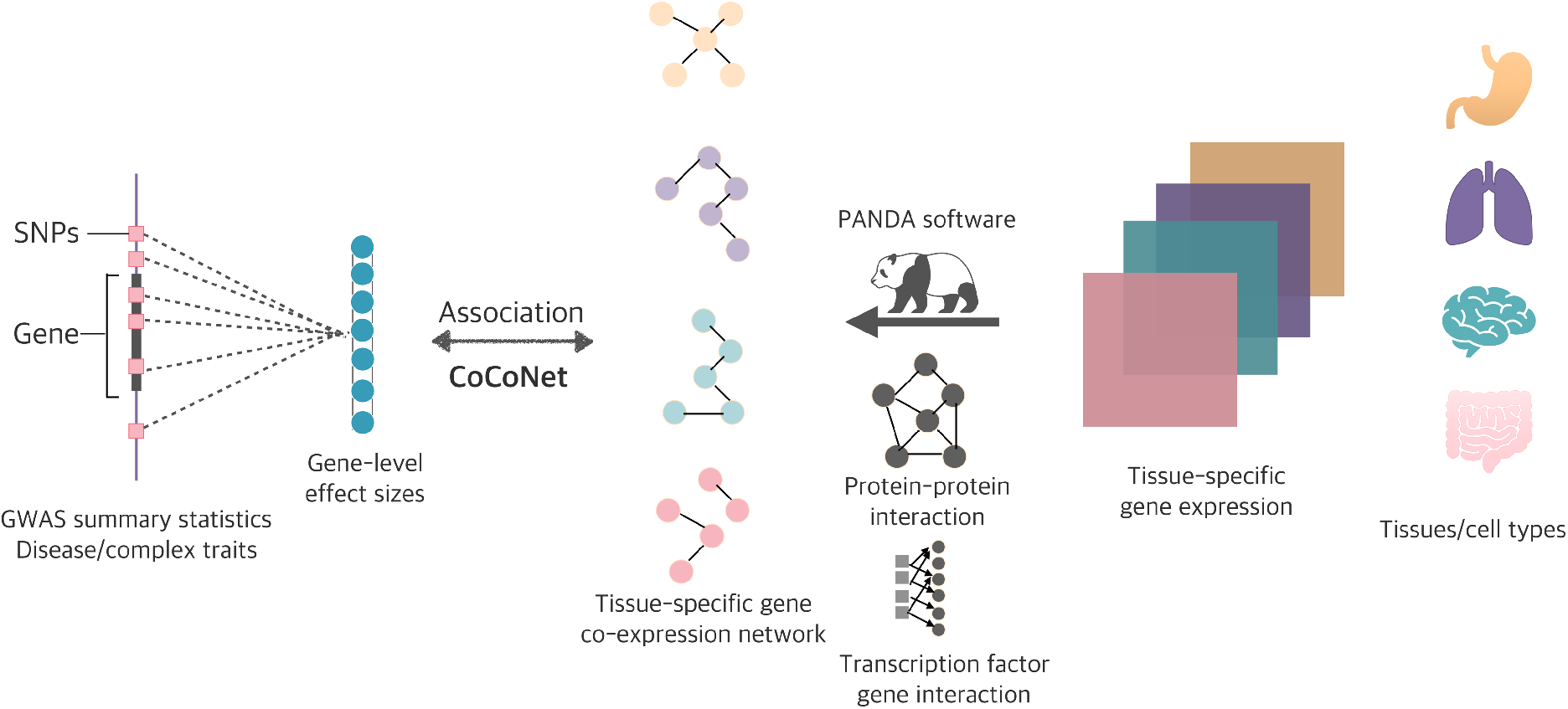
Schematic of CoCoNet for identifying trait relevant tissue/cell type using information from gene co-expression networks. From left to middle: we calculate gene level effect sizes from GWAS summary statistics for the trait of interest and treat them as an *m* dimensional vector of outcomes. From right to middle: we infer the tissue specific gene co-expression network for each tissue/cell type using the PANDA software, which requires the tissue-specific gene expression matrix, existing protein-protein interaction network information, as well as existing transcription factor and gene binding information as input. For the trait of interest, we examine one tissue at a time and we model the gene-level effect sizes for the trait as a function of the gene co-expression matrix using a covariance regression network model. We infer parameters in the model through composite likelihood. We calculate the maximum composite likelihood for each tissue and eventually rank tissues by the corresponding log likelihoods.

**Fig 2.**
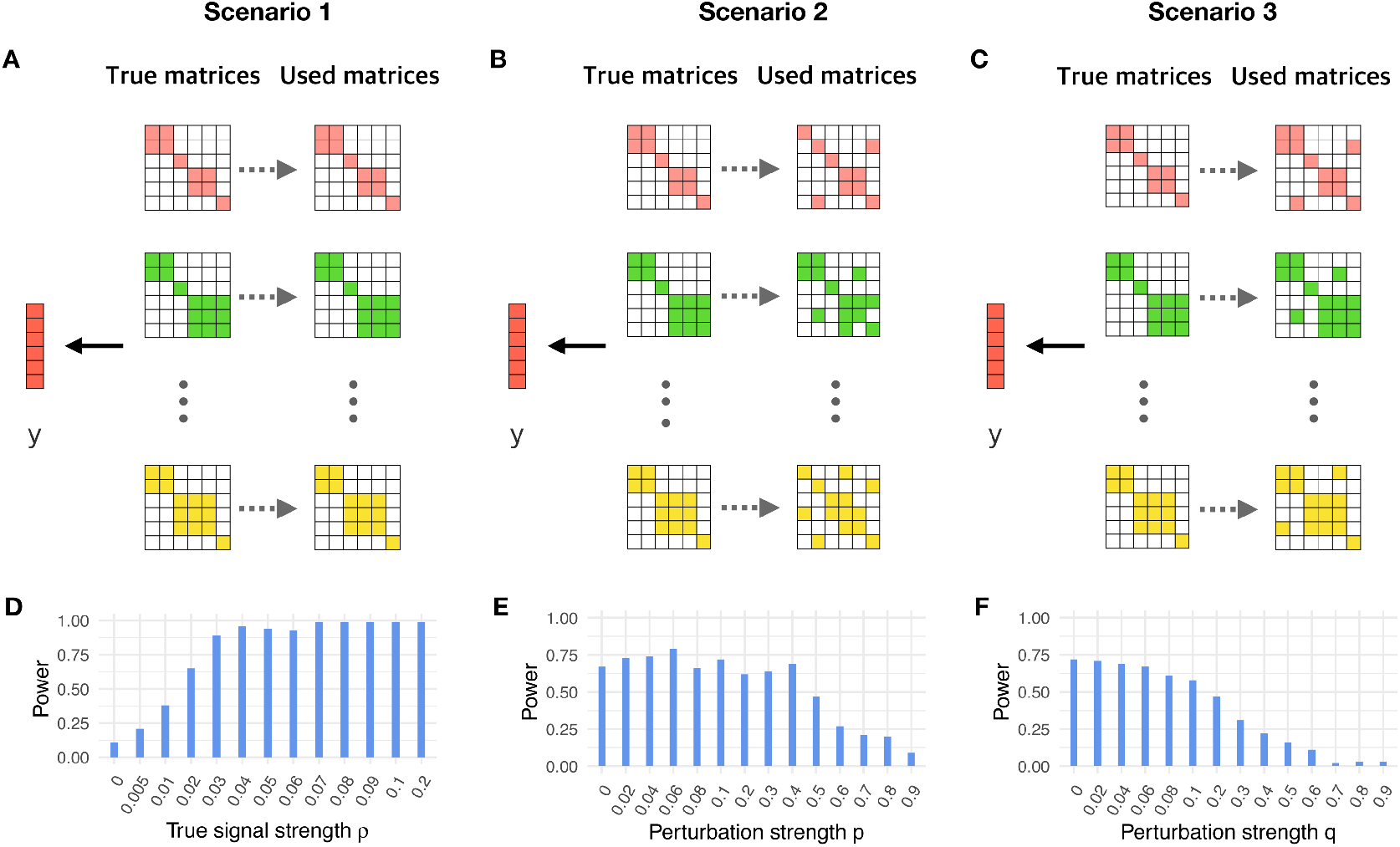
Power of CoCoNet to identify trait relevant tissues in simulations. (A): In scenario I, we selected one of the ten tissues as the trait-relevant tissue (second row) and simulated gene-level effect sizes based on the adjacency matrix (first row). We treated the adjacency matrices all as observed (third row) and fit the model using each of the ten matrices. (B): In scenario II, the data are generated in the same way as in scenario I but we fit the model using tissue-specific matrices that are noisy versions of the truth (third row). In particular, we assumed that each connected gene pair in the true adjacency matrices has a probability of *p* being un-observed, and each unconnected gene pair has a probability *q* to be falsely assigned as connected. (C): In scenario III, the data are again generated in the same way as in scenario I but we fit the model using tissue-specific matrices that are noisy versions of the truth (third row). In particular, we randomly converted a proportion *q* of unconnected gene pairs in the true adjacency matrix to be connected. (D): The power of CoCoNet for identifying the correct tissue (y-axis) increases with increasing signal strength measured by 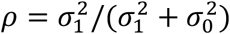 (x-axis) in scenario I. (E): The power of CoCoNet for identifying the correct tissue (y-axis) gradually decreases with increasing noise level characterized by *p* (x-axis) in scenario II. (F): The power of CoCoNet for identifying the correct tissue (y-axis) gradually decreases with increasing noise level characterized by *q* (x-axis) in scenario III.

In scenario II, when we fit the simulated data, we assumed that we did not observe the true tissue-specific gene co-expression adjacency matrices. Instead, we only observed a noisy version of them (Fig 2B). In particular, we added noise into each of the adjacency matrices by assuming that each pair of the connected genes has a probability of *p* being unobserved, and each pair of unconnected genes has a probability *q* to be falsely assigned as connected. For each fixed value *p*, we calculated *q* such that in expectation the proportion of connected gene pairs in the adjacency matrix equals to that of the original one. Here, we fixed *ρ* = 0.02 and varied *p* to examine its influence on parameter estimates and power. As expected, with increasing *p*, the observed adjacency matrix deviates further from the true networks. Subsequently the accuracy of the parameter estimates *ρ* (S2 Fig B and E) and 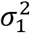 (S4 Fig B and E) reduces; though the 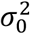 estimates are relatively stable and accurate (S3 Fig B and E). Importantly, however, the power of our method in detecting the trait relevant tissue was relatively stable for small *p* (e.g. *p* < 0.4), remaining around 0.7 (Fig 2E). Certainly, with increasingly large *p*, the power of our method gradually decreases towards the null expectation of 0.1.

In scenario III, we also assumed that the observed gene co-expression adjacency matrices are a noisy version of the truth (Fig 2C). Different from scenario II, however, we assumed here that the true adjacency matrices can be considered as a subset of the observed adjacency matrices, mimicking the setting where only the regulatory network of a set of core genes influences gene-level effect sizes on the trait of interest. In particular, we assigned noise to the networks by randomly converting unconnected gene pairs in the true adjacency matrices to be connected with a probability *q*. Here, we fixed *ρ* = 0.02 and varied the value of *q* to examine its influence on parameter estimation and power. As expected, because the noise added to the matrices makes it hard to perform accurate estimation, the accuracy of *ρ* (S2 Fig C and F) and 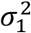 (S4 Fig C and F) become worse with increasing *q*; though the 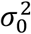 estimates are relatively stable and accurate (S3 Fig C and F). Importantly, however, the power of our method in detecting trait relevant tissues remains stable and is around 0.7 across a range of reasonable *q* (*q*<0.1); the power only starts to decrease with continuously increasing *q* (Fig 2F).

#### Real data application: inferring trait-relevant tissues with bulk RNAseq

We focus our first real data application on eight different disease GWASs that include four neurological disorders and four autoimmune disorders. We focus on these diseases because their disease relevant tissues have been reasonably well characterized and the corresponding GWASs have sufficiently large sample sizes (>12,000). Here, we aim to identify for each disease the trait-relevant tissue among 38 tissues obtained from GTEx. To do so, for each of the 38 tissues in turn, we followed Sonawane et al. [45] and used the PANDA software to infer a tissue-specific gene co-expression adjacency matrix. These inferred adjacency matrices contain tissue specific information (S5 Fig and S6 Fig). For example, the adjacency matrices for the three brain tissues (basal ganglia, cerebellum, and brain other) are all clustered together. Similarly, the adjacency matrices for the intestinal tissues (stomach, colon-transverse, intestine terminal ileum, and colon sigmoid) are all clustered together. With the adjacency matrices, we applied CoCoNet to examine one tissue at a time and ranked the relevance of tissue to disease by log likelihood.

Overall, we found that the top relevant tissues identified for neurological disorders are generally brain tissues and the top relevant tissues identified for autoimmune disorders are generally intestinal tissues (Table 1 and S7 Fig). For example, at least one brain tissue is identified either as the most relevant or the second most relevant tissue for all four neurological disorders (Table 1 and S7 Fig A-D). Similarly, at least one intestinal tissue is identified either as the most relevant or the second most relevant tissue for all four autoimmune disorders (Table 1 and S7 Fig E-H). The inferred ranking of tissues for each disease does not depend on the sample size of each tissue. Specifically, Spearman’s rank correlation between tissue ranking and tissue sample size ranges from −0.32 to 0.13 for the eight diseases, with none of them being significant (S8 Fig). Besides CoCoNet, we analyzed the same data with RolyPoly and LDSC, both of which use tissue-specific gene expression information in place of tissue-specific gene co-expression network information for trait-tissue relevance inference. In the comparison, we found that the ranking of brain tissues for neurological disorders obtained by CoCoNet is higher than that obtained by RolyPoly or LDSC (Fig 3A). Similarly, the ranking of colon tissues for autoimmune disorders obtained by CoCoNet is also higher than that obtained by RolyPoly or LDSC (Fig 3B). The comparative results between CoCoNet and RolyPoly/LDSC suggests that tissue-specific gene co-expression network provides valuable trait-tissue relevance information, more so than the information provided by tissue-specific gene expression pattern used in RolyPoly or LDSC.

**Fig 3.**
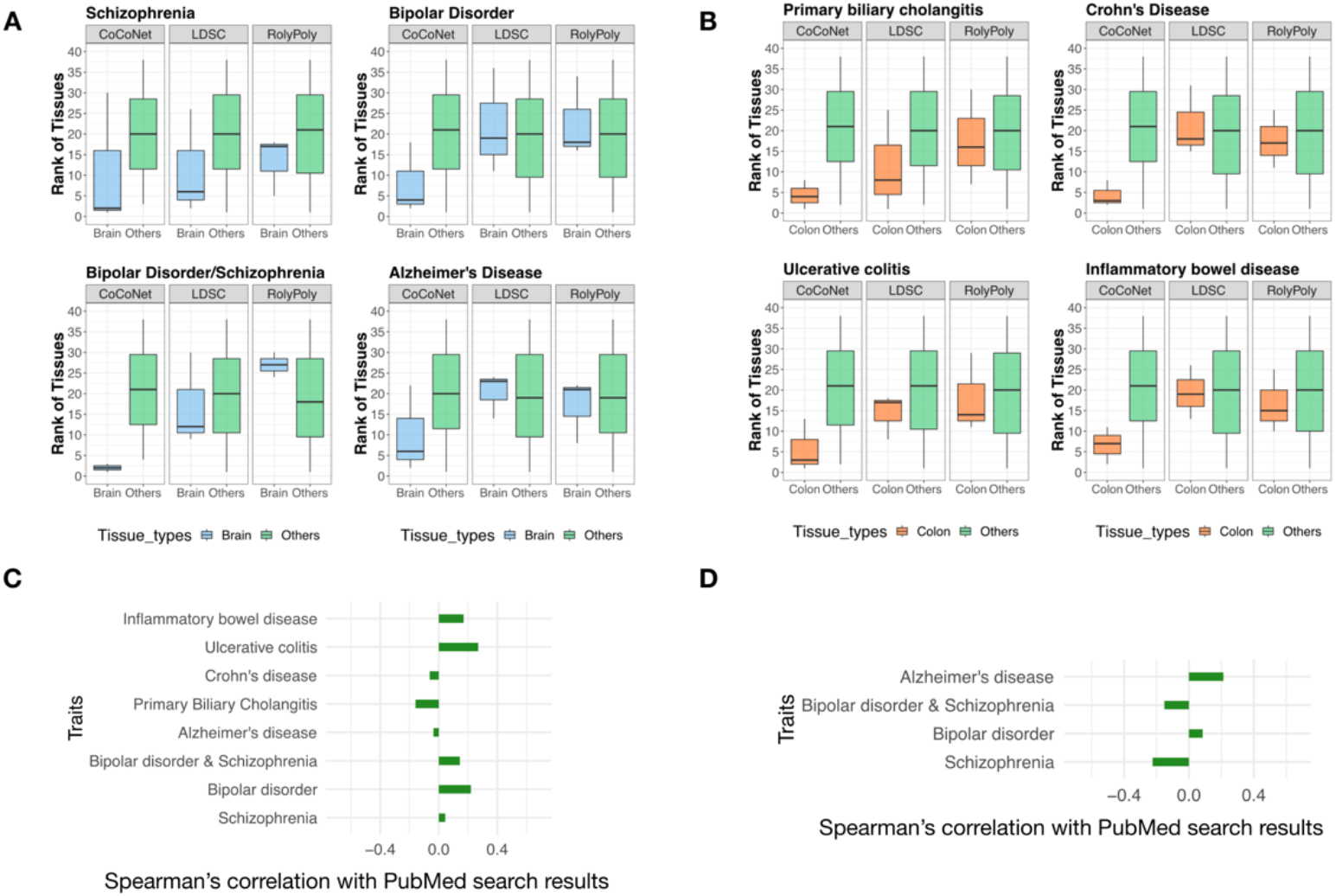
Comparison of tissue rankings by CoCoNet, LDSC, and RolyPoly for eight GWAS traits. (A): For each of the four neurological diseases, we plotted the rank (y-axis) of three brain tissues (colored blue; including brain cerebellum, brain basal ganglia, and brain other) and the rank of the remaining 35 tissues (colored green) in separate boxplots. The rank of brain tissues obtained using CoCoNet is often lower than that of the non-brain tissues (first column in each panel). The rank difference between the two types of tissues obtained by CoCoNet (first column in each panel) is often more pronounced than that obtained by either LDSC (second column) or RolyPoly (third column) for the four neurological diseases. (B): For each of the four autoimmune diseases, we plotted the rank (y-axis) of three colon tissues (colored orange; including colon sigmoid, colon transverse, and intestine terminal ileum) and the rank of the remaining 35 tissues (colored green) in separate boxplots. The rank of colon tissues obtained using CoCoNet is often lower than that of the non-colon tissues (first column in each panel). The rank difference between the two types of tissues obtained by CoCoNet (first column in each panel) is often more pronounced than that obtained by either LDSC (second column) or RolyPoly (third column) for the four autoimmune diseases. (C): We compared the ranking of tissues obtained by CoCoNet with that obtained by PubMed search for 8 GWAS traits, in the bulk RNAseq data. The CoCoNet and PubMed search results are positively correlated with each other for five traits. (D): We compared the ranking of cell types obtained by CoCoNet with that obtained by PubMed search for 4 GWAS traits, in the single cell RNAseq data. The CoCoNet and PubMed search results are positively correlated with each other for two traits.

**Table 1.** Top five tissue types identified among the 38 tissues in the GTEx bulk RNAseq data for each of the eight GWAS traits.

| Traits | Top1 | Top2 | Top3 | Top4 | Top5 |
| --- | --- | --- | --- | --- | --- |
| Schizophrenia | Brain basal ganglia | Brain other | Uterus | Skin | Tibial Nerve |
| Bipolar Disorder | Prostate | Brain basal ganglia | Esophagus mucosa | Breast | Brain cerebellum |
| Bipolar Disorder/<br>Schizophrenia | Brain basal ganglia | Brain cerebellum | Spleen | Brain other | Thyroid |
| Alzheimer's Disease | Testis | Brain other | Colon sigmoid | Colon transverse | Intestine terminal ileum |
| Primary biliary cholangitis | Intestine terminal ileum | Esophagus mucosa | Colon sigmoid | Stomach | Artery coronary |
| Crohn's Disease | Testis | Colon transverse | Artery coronary | Colon sigmoid | Heart atrial appendage |
| Ulcerative colitis | Stomach | Whole blood | Colon sigmoid | Fibroblast cell line | Breast |
| Inflammatory bowel disease | Testis | Colon sigmoid | Heart atrial appendage | Skeletal muscle | Tibial Nerve |

Next, we attempted to validate the identified trait-relevant tissue by performing a PubMed search following the main idea in [8]. Specifically, we reasoned that, if a tissue is relevant to the disease of interest, then there would be previous publications studying the disease on the particular tissue. Therefore, by counting the number of previous publications using the key word pairs of trait and tissue, we would have a reasonable quantitative estimate on the relevance between the trait and the corresponding tissue, which can serve to validate the results obtained by our method. Here, we followed such intuition and counted the number of previous publications on pairs of trait and tissue (details in Materials and Methods). For each trait in turn, we ranked tissues by the number of previous publications on the trait-tissue pair. We then calculated the Spearman’s correlations between the tissue rank obtained by CoCoNet and the tissue rank obtained by PubMed search (Fig 3D). Overall, the tissue ranking results obtained by CoCoNet is reasonably consistent with PubMed search results for majority of traits: the Spearman’s rank correlation between these two approaches is 0.07 on average (median = 0.09), with ranges from −0.15 (for PBC) to 0.26 (for UC); the correlation is positive for five out of eight traits. The correlation is lower than those obtained using histone modification marks (median = 0.417 based on [8]), consistent with the previous observation that histone modification contains more information than gene expression data for trait-relevant tissue inference [44].

As a concrete example, for schizophrenia, our method identified the basal ganglia as the most relevant tissue, which is consistent with PubMed search results. Basal ganglia is a functional brain region involved in a variety of activities including motor movements, cognition and emotion. Basal ganglia contains most of the dopamine neurons in the brain and dopamine is the first neurotransmitter implicated for schizophrenia [46]. Indeed, the dopaminergic system of the basal ganglia is known to display several anomalies in schizophrenia and is itself a target of various antipsychotic drugs [47, 48]. As another example, for Crohn’s disease, our method identified colon transverse and colon sigmoid as the most relevant tissues. The results are consistent with PubMed search results, which identified colon transverse and colon sigmoid as the most relevant tissues. Crohn’s disease is a chronic inflammatory disease that is known to affect all segments of the gastrointestinal tract, with particular pathological influence on the terminal ileum and colon [49].

#### Real data application: inferring trait-relevant cell types with single cell RNAseq

Our second application focuses on GWASs of the four neurological diseases described in the previous section. For each disease in turn, we aimed to identify the trait-relevant cell types. To do so, we constructed gene co-expression adjacency matrices for each of the 10 cell types from two donors using single cell sequencing data [41]. The 10 cell types include five neuronal cell types, four glial cell types, and one endothelial cell type. For each cell type in turn, we inferred a gene co-expression adjacency matrix through the same analytic pipeline described in the previous section. These inferred gene co-expression adjacency matrices contain cell type specific information (S9 Fig and S10 Fig). For example, astrocytes from two donors were clustered together, along with other glia cells such as microglia, oligodendrocytes, and oligodendrocyte precursor cells. The neuronal cells, such as glutamatergic neurons, GABAergic interneurons, pyramidal neurons, and granule neurons were also clustered together. Therefore, in our following analysis, we combined the adjacency matrices for the same cell type from the two donors into a single adjacency matrix through union operation. For each disease in turn, we applied our method to examine one cell type at a time and ranked the relevance of cell types to the disease by log likelihood (Table 2). The ranking of cell types for each neurological disease does not depend on the number of cells in each cell type (Spearman’s rank correlation ranges from −0.39 to 0.33; only one statistically significant) (S11 Fig). Finally, we also compared the tissue ranking results obtained by CoCoNet to those obtained through PubMed search (Fig 3D). Likely due to high experimental noise in single cell RNAseq data, the cell ranking results obtained by CoCoNet is consistent with PubMed search results for two out of four diseases: the rank correlation is 0.21 for Alzheimer’s disease and 0.08 for Bipolar disorder.

**Table 2.** Top five cell types identified among 10 cell types in the single cell RNAseq data for each of the four neurological disorders.

| Traits | Top1 | Top2 | Top3 | Top4 | Top5 |
| --- | --- | --- | --- | --- | --- |
| Schizophrenia | Granule neurons | Microglia | Endothelial cell | Oligodendrocyte precursor cells | Pyramidal neurons |
| Bipolar Disorder | Pyramidal neurons | Microglia | Oligodendrocyte precursor cells | Oligodendrocyte | Astrocytes |
| Bipolar Disorder/<br>Schizophrenia | Oligodendrocytes | Endothelial cell | Glutamatergic neurons | Astrocytes | Granule neurons |
| Alzheimer's Disease | GABAergic interneurons | Oligodendrocyte precursor cells | Astrocytes | Microglia | Endothelial cell |

Cell types identified in the top trait-tissue relevance list for these neurological diseases often make biological sense (S12 Fig). For example, for Alzheimer’s disease, we ranked GABAergic interneurons, oligodendrocyte precursor cells, astrocytes, and microglia as top relevant cell types. Both astrocytes and microglia were identified by PubMed search as the most relevant cell types. These two glia cell types are activated by the pathological peptide amyloid beta and release proinflammatory mediators to induce neuronal death [50, 51]. Similarly, significant reductions in GABA levels and reduced GABAergic innervations are commonly observed during the progression of Alzheimer’s disease. Such GABA reduction leads to eventual disruption of the neuronal circuitry and impaired cognition [52]. Finally, vulnerability of oligodendrocyte precursor cells under Alzheimer’s pathology can induce myelin breakdown and loss of myelin sheath, which characterizes the earliest stage of Alzheimer’s disease [53]. As another example, for bipolar disorder, both pyramidal neurons and various glia cells are selected as top relevant cell types. In patients with bipolar disorder, the size of pyramidal neurons in the CA1 region of the hippocampus are often decreased, suggesting pathogenic effects of bipolar disorder on pyramidal neurons [54]. Similarly, it has been shown that decreased density of glial cells may contribute to the pathological changes observed in neurons in bipolar disorder patients [55].

## Discussion

We have presented a new method to leverage gene co-expression patterns for inferring trait-relevant tissue or cell types. With the real data examples, we show that tissue-specific gene coexpression patterns contain valuable information for inferring trait-tissue relevance. Our results support the hypothesis in the omnigenic model that tissue specific gene co-expression networks underlie disease etiology.

There are two important ingredients in our method. The first key ingredient is the tissue-specific gene co-expression matrix. In the present study, we have primarily used the adjacency matrix based on hard thresholding the tissue-specific gene co-expression matrices obtained from [45]. Because of the small sample sizes relative to the number of genes in the gene expression study, matrix sparsity introduced by hard thresholding is crucial for improving the signal contained in the coexpression matrices. While hard thresholding is one of the easiest approaches to introduce matrix sparsity, other sophisticated sparse matrix methods, such as graphical lasso [56, 57], may have added benefits and are worth future exploration. In addition, we have primarily focused on using the adjacency matrix directly in the covariance function. The covariance function used in CoCoNet can be general and can consist of higher order terms of the adjacency matrix through the polynomial matrix construction. We have implemented our composite likelihood method to accommodate high order terms of the adjacency matrix as determined by the parameter *K*. However, we found that models with larger *K* tend to have higher BIC in the two data applications we examined here, indicating that direct connections between genes may contain sufficient information for trait tissue relevance inference. Besides the use of polynomial matrix construction, alternative ways to construct the covariance function, such as the use of graph Laplacian matrix [58], could be an interesting area for future exploration. We also note that our adjacency matrices are constructed based on the PANDA software. The PANDA software can integrate multiple data sources to facilitate the construction of tissue-specific gene co-expression matrices and are thus commonly used previously. The data sources used in PANDA include gene expression data, protein-protein interaction, as well as transcription factor binding information. Besides the default PANDA data sources, there are many other software and informative databases for constructing gene-gene co-expression networks. For example, GIANT (Genome-scale Integrated Analysis of gene Networks in Tissues) is a tissue-specific interaction network database that integrates physical interaction, co-expression, miRNA binding motif and transcription factor binding site data [59]. Such a database can be paired with PANDA to facilitate accurate gene co-expression network construction. Besides PANDA, various other methods for inferring tissue-specific gene coexpression patterns can also be used in pair with CoCoNet. Some of these methods, such as ARACNE (Algorithm for the Reconstruction of Accurate Cellular Networks) [60] and WGCNA (weighted gene co-expression network analysis) [61], only rely on gene expression data. Some of these methods, such as GRAM (Genetic Regulatory Modules) [62], CLT (Context likelihood of relatedness) [63] and GENIE3 (GEne Network Inference with Ensemble of trees) [64], can integrate gene expression data together with transcription factor information. While none of these methods can incorporate protein-protein interaction information as PANDA does, exploring the benefits of different methods in constructing tissue-specific gene co-expressing pattern for trait-tissue relevance inference remains an important future area of research.

The second key ingredient of our method is the gene-level effect sizes for the GWAS trait. In the present study, we have primarily used gene-level per-SNP heritability estimates as the gene-level effect sizes. The gene-level heritability not only has a natural genetic interpretation but is also directly linked to the gene-level effect size in the sequence kernel association test (SKAT) [65], which has been widely used for SNP set tests. Besides SKAT, many SNP set methods exist for carrying out gene-level tests. Common SNP set methods include the versatile gene-based test (VEGAS) [66], the MinP approach [67], standard combination statistics (e.g. Fisher, Sidat, and Simes) [68], and the principal component analysis regressions [69]. Different SNP set methods have different benefits and disadvantages [67, 70, 71], and consequently, gene-level effect sizes constructed from different methods may contain different levels of information for inferring trait-tissue relevance. CoCoNet can be paired with gene-level statistics constructed from any SNP set test and it would be important to explore the benefits of different gene-level statistics for trait-tissue relevance inference in the future.

Finally, we have only focused on ranking tissues for a given disease instead of directly testing the tissue relevance to the disease. Ranking tissues requires model selection, which is an easier analysis task than carrying out hypothesis testing for tissue relevance to disease. Hypothesis testing on the tissue relevance to disease using gene co-expression patterns is challenging because of the strong correlation among gene co-expression patterns constructed from different tissues. Specifically, the gene co-expression matrices constructed in different tissues are highly correlated with each other. Because of the high correlation across gene co-expression matrices, the estimated maximum likelihood across different tissues using CoCoNet can be similar to each other. Subsequently, the naïve p-value for testing trait-tissue relevance would become significant even for trait-irrelevant tissues, simply due to tissue-tissue correlation. Therefore, a naïve hypothesis test would lead to an undesirably large number of false positives. Previous studies using gene-level epigenomic annotations for trait-tissue relevance inference have also encountered a similar phenomenon. Indeed, previous studies have to either introduce a large set of covariates to account for tissue-tissue correlation [34], or rely on a mixture model to formulate the hypothesis test into a model selection framework [8]. Unfortunately, we found that adjusting for known gene-level covariates is not effective for addressing correlation among gene co-expression matrices across tissues. In addition, a mixture model-based model selection approach is technically challenging in our setting due to the small difference in the estimated maximum likelihood across tissues. Therefore, we have resorted to the easier task of ranking tissues for a given disease and focus on the top selected tissues for biological interpretation. Future methodological innovations are needed to directly infer statistical confidence in the ranking itself or to develop effective hypothesis tests to incorporate gene co-expression pattern for trait-relevant tissue inference.

## Acknowledgements

This study was supported by the National Institutes of Health (NIH) grants R01HG009124 and R01GM126553, and the National Science Foundation (NSF) grant DMS1712933. This project has been made possible in part by grant number 2018-181314 from the Chan Zuckerberg Initiative DAF, an advised fund of Silicon Valley Community Foundation. LS was also partially supported by NIH grant R01HL133221 (PI Smith). The funders had no role in study design, data collection and analysis, decision to publish, or preparation of the manuscript.

