## Supplementary materials for "Leveraging Gene Co-expression Patterns to Infer Trait-Relevant Tissues in Genome-wide Association Studies"

**Supplementary Text**

**CoCoNet Inference**

Our composite likelihood for the covariance regression network model is defined based on the product of pair-wise likelihood. For each pair of genes $i$ and gene $j$, we have:

$$\left( \begin{matrix} y_{i} \\ y_{j} \end{matrix} \right)\sim BN(\left( \begin{matrix} \mu\\ \mu\end{matrix} \right),\sum_{k=0}^{K} \sigma_{k}^{2}\left( \begin{matrix} a_{ii}^{\left( k \right)} & a_{ij}^{\left( k \right)} \\ a_{ij}^{\left( k \right)} & a_{jj}^{\left( k \right)} \end{matrix} \right))$$

where ***A*** is the adjacency matrix, and $\boldsymbol{A}^{k}=(a_{ij}^{\left( k \right)})$ is the *k-*th power of $\boldsymbol{A}$, for any integer *k*.

We can write $\sum_{k=0}^{K} \sigma_{k}^{2}\boldsymbol{A}^{k}=\left( \begin{matrix} b_{ii} & b_{ij} \\ b_{ji} & b_{jj} \end{matrix} \right)$,

where

$$b_{ii}=b_{jj}=\sigma_{0}^{2}+\sigma_{1}^{2}a_{ii}^{(1)}+ \sigma_{2}^{2}a_{ii}^{(2)}+\ldots\sigma_{K}^{2}a_{ii}^{\left( K \right)}=\sigma_{0}^{2}$$

since we set $a_{ii}^{\left( k \right)}$ to be zero for $k\geq1$,

and

$b_{ij}=b_{ji}=\sigma_{1}^{2}a_{ij}^{(1)}+ \sigma_{2}^{2}a_{ij}^{(2)}+\ldots\sigma_{K}^{2}a_{ij}^{(K)}$.

The likelihood function for a pair of gene $i$ and $j$:

$$\log P\left( (y_{i},y_{j}|\mu,\sigma_{k}^{2}) \right)=-\frac{1}{2}\log\left( b_{ii}^{2}-b_{ij}^{2} \right)-\frac{1}{2\left( b_{ii}^{2}-b_{ij}^{2} \right)}\left( \begin{matrix} y_{i}-\mu\\ y_{j}-\mu\end{matrix} \right)^{T}\left( \begin{matrix} b_{jj} & -b_{ij} \\ -b_{ji} & b_{ii} \end{matrix} \right)\left( \begin{matrix} y_{i}-\mu\\ y_{j}-\mu\end{matrix} \right)-\frac{1}{2}\log\left( \frac{1}{2\pi} \right)=-\frac{1}{2}\log\left( b_{ii}^{2}-b_{ij}^{2} \right)-\frac{1}{2\left( b_{ii}^{2}-b_{ij}^{2} \right)}[\left( \left( y_{i}-\mu\right)^{2}+\left( y_{j}-\mu\right)^{2} \right)b_{ii}-2b_{ij}\left( y_{i}-\mu\right)\left( y_{j}-\mu\right)]-\frac{1}{2}\log\left( \frac{1}{2\pi} \right)$$

Full composite likelihood function:

$$l\left( \theta\right)=\sum_{i=1}^{m}\Sigma_{j>i}^{m}logP(y_{i},y_{j}|\mu,\sigma_{k}^{2})$$

$$\log P=\frac{1}{2\left( m-1 \right)}\sum_{i=1}^{m} \sum_{j\neq i} logP(y_{i},y_{j}|\mu,\sigma_{k}^{2})=\frac{1}{2(m-1)}\sum_{i=1}^{n} \sum_{j\neq i} [-\frac{1}{2}\log\left( b_{ii}^{2}-b_{ij}^{2} \right)-\frac{1}{2\left( b_{ii}^{2}-b_{ij}^{2} \right)}[\left( \left( y_{i}-\mu\right)^{2}+\left( y_{j}-\mu\right)^{2} \right)b_{ii}-2b_{ij}\left( y_{i}-\mu\right)\left( y_{j}-\mu\right)]-\frac{1}{2}\log\left( \frac{1}{2\pi} \right)]$$

**Supplementary Figures and Tables**

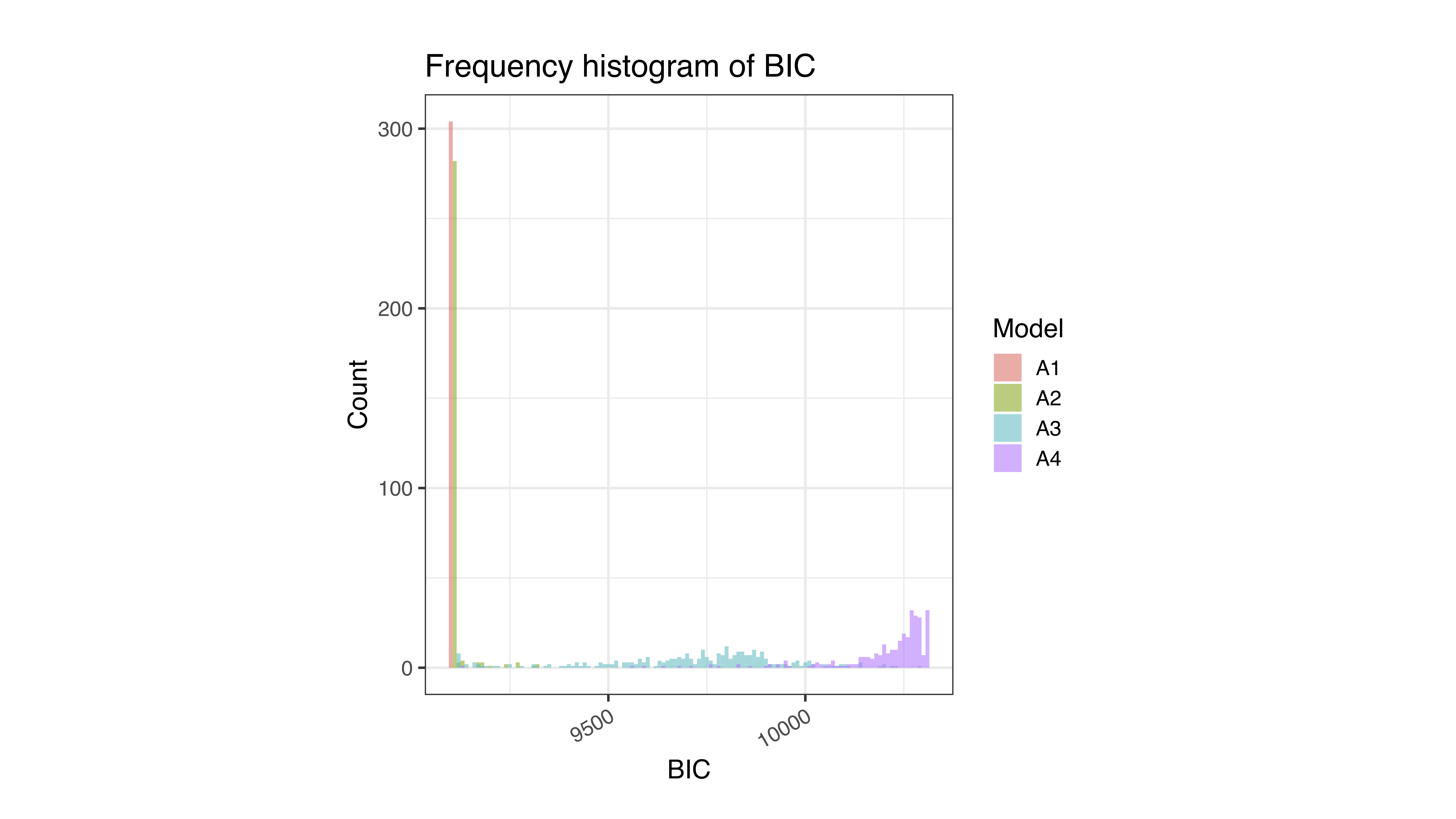

**S1 Fig. A CoCoNet model with a small *K* is often preferred than a CoCoNet model with a large *K* in real data sets.** Here, *K* is the number of matrices included in the covariance function. In the real data application, we analyzed all pairs of 38 GTEx tissues and 8 GWAS traits. For each of the 304 trait-tissue pairs, we fit four different CoCoNet models with K ranging from 1 to 4. The above histogram shows the Bayesian information criterion (BIC) values (x-axis) across all these CoCoNet models. Models with different *K* are colored differently: A1 represents a model with 1^st^ power of the adjacency matrix (red); A2 represents a model with both 1^st^ and 2^nd^ power of the adjacency matrix (green); A3 represents a model with up to the 3^rd^ power of the adjacency matrix (blue); A4 represents a model with up to the 4^th^ power of the adjacency matrix (purple). The results suggest that a model with a low value of *K* (1 or 2) often comes with a lower BIC and is thus often preferred than a model with a high *K*.

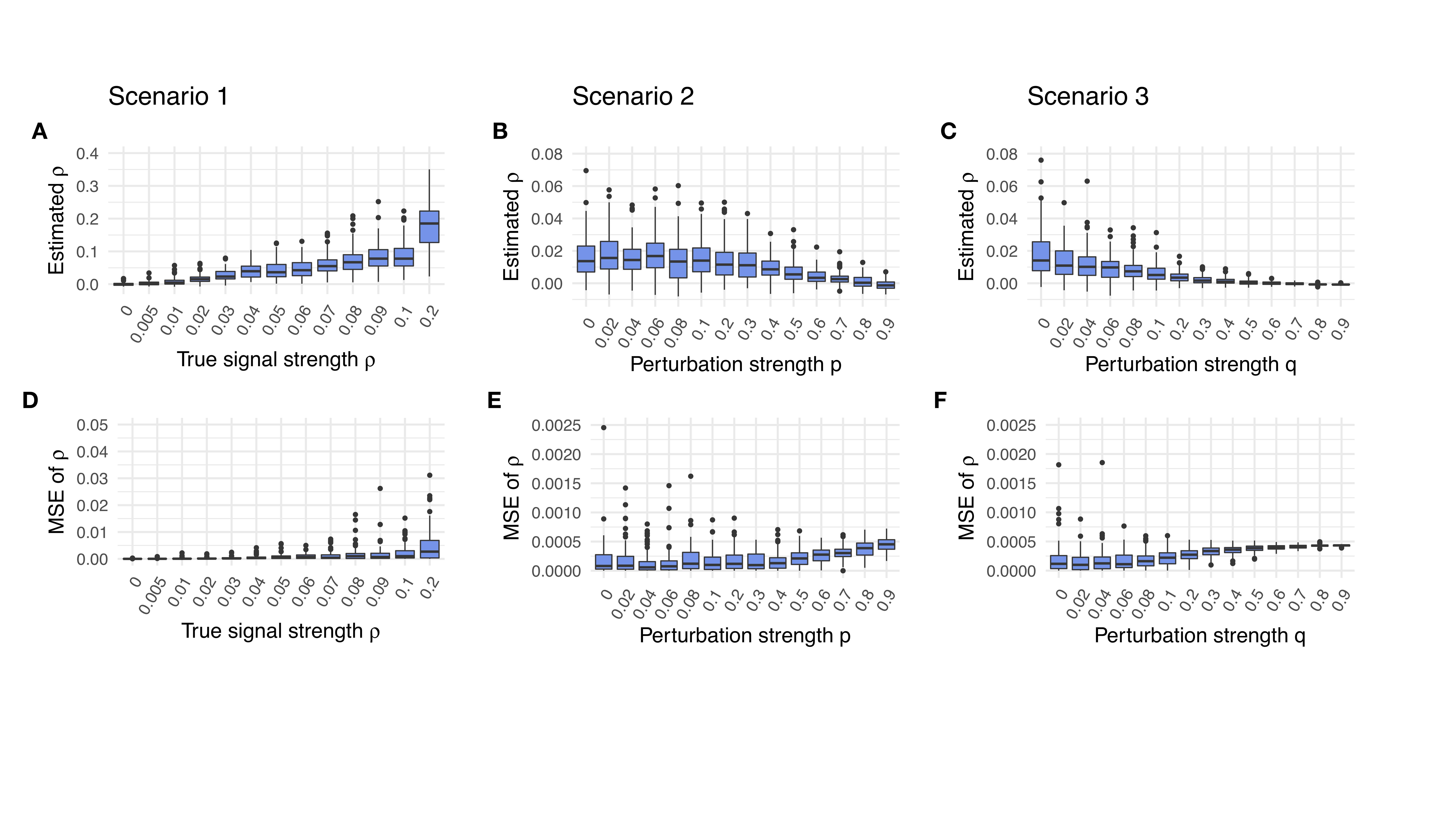

**S2 Fig. Parameter estimation of the signal strength parameter** $\boldsymbol{\rho}$ **in simulations.** (A): Boxplot shows the $\rho$ estimates across 100 simulation replicates for each true $\rho$ (x-axis) in the simulation scenario I. (B): Boxplot shows the $\rho$ estimates across 100 simulation replicates for each parameter *p* (x-axis) in the simulation scenario II. Here, true $\rho=0.02$. Increasing parameter *p* adds increasingly large noise to the tissue-specific adjacency matrices, thus leading to downward biased estimation of $\rho$. (C): Boxplot shows the $\rho$ estimates across 100 simulation replicates for each parameter *q* (x-axis) in the simulation scenario III. Here, true $\rho=0.02$. Increasing parameter *q* adds increasingly more noise to the tissue-specific adjacency matrices, thus leading to downward biased estimation of $\rho$. (D-F): mean squared error (MSE; y-axis) measures the accuracy of $\rho$ estimates across the three simulation scenarios.

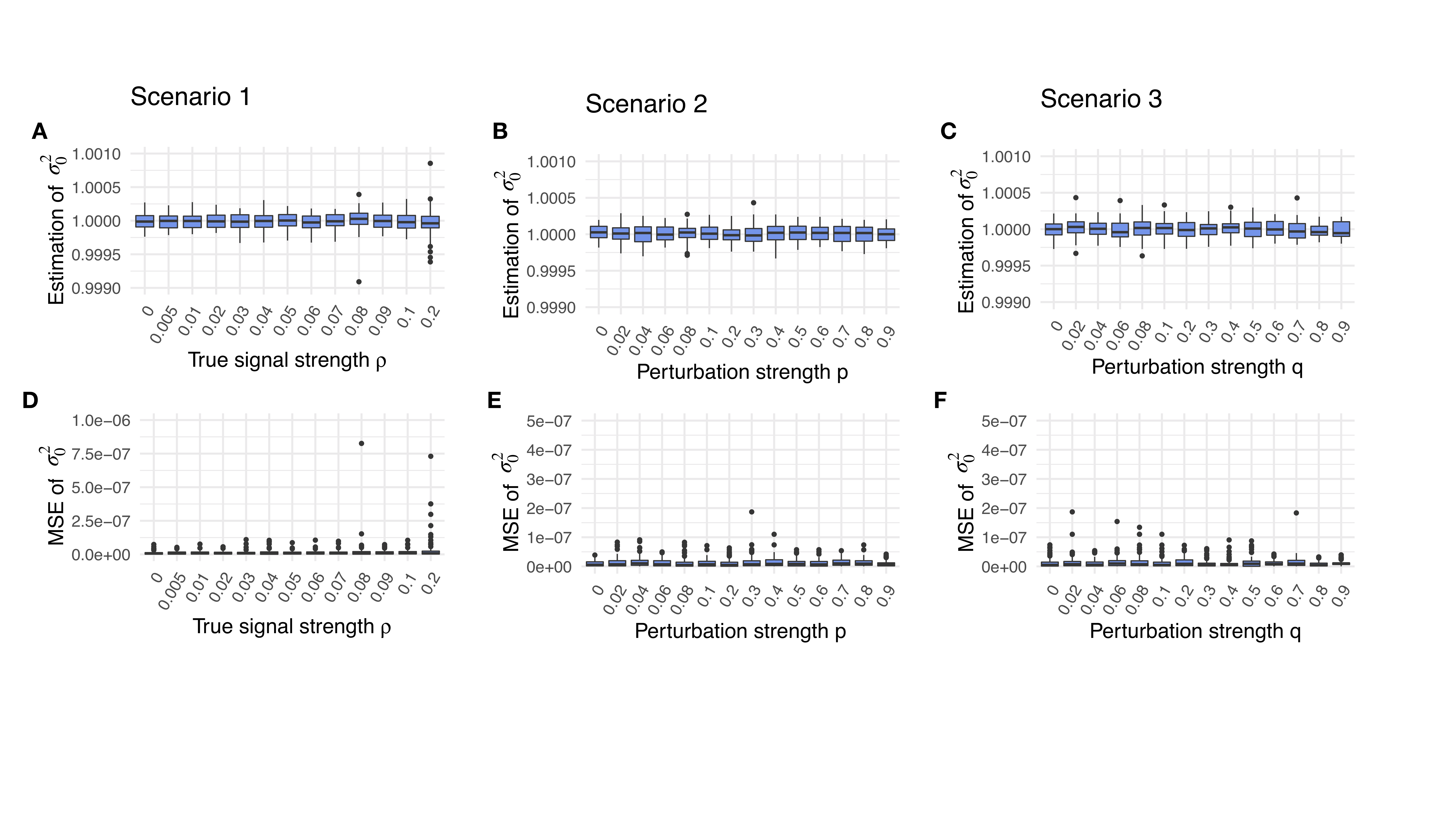

**S3 Fig. Parameter estimation of the parameter** $\boldsymbol{\sigma}_{\boldsymbol{0}}^{\boldsymbol{2}}$ **in simulations.** (A): Boxplot shows the $\sigma_{0}^{2}$ estimates across 100 simulation replicates for each signal strength parameter $\rho$ (x-axis) in the simulation scenario I. (B): Boxplot shows the $\sigma_{0}^{2}$ estimates across 100 simulation replicates for each parameter *p* (x-axis) in the simulation scenario II. Here, true $\sigma_{0}^{2}=0.98$. Increasing parameter *p* adds increasingly large noise to the tissue-specific adjacency matrices, but does not appear to strongly influence the estimation of $\sigma_{0}^{2}$. (C): Boxplot shows the $\sigma_{0}^{2}$ estimates across 100 simulation replicates for each parameter *q* (x-axis) in the simulation scenario III. Here, true $\sigma_{0}^{2}=0.98$. Increasing parameter *q* adds increasingly large noise to the tissue-specific adjacency matrices, but does not appear to strongly influence the estimation of $\sigma_{0}^{2}$. (D-F): mean squared error (MSE; y-axis) measures the accuracy of $\sigma_{0}^{2}$ estimates across the three simulation scenarios.

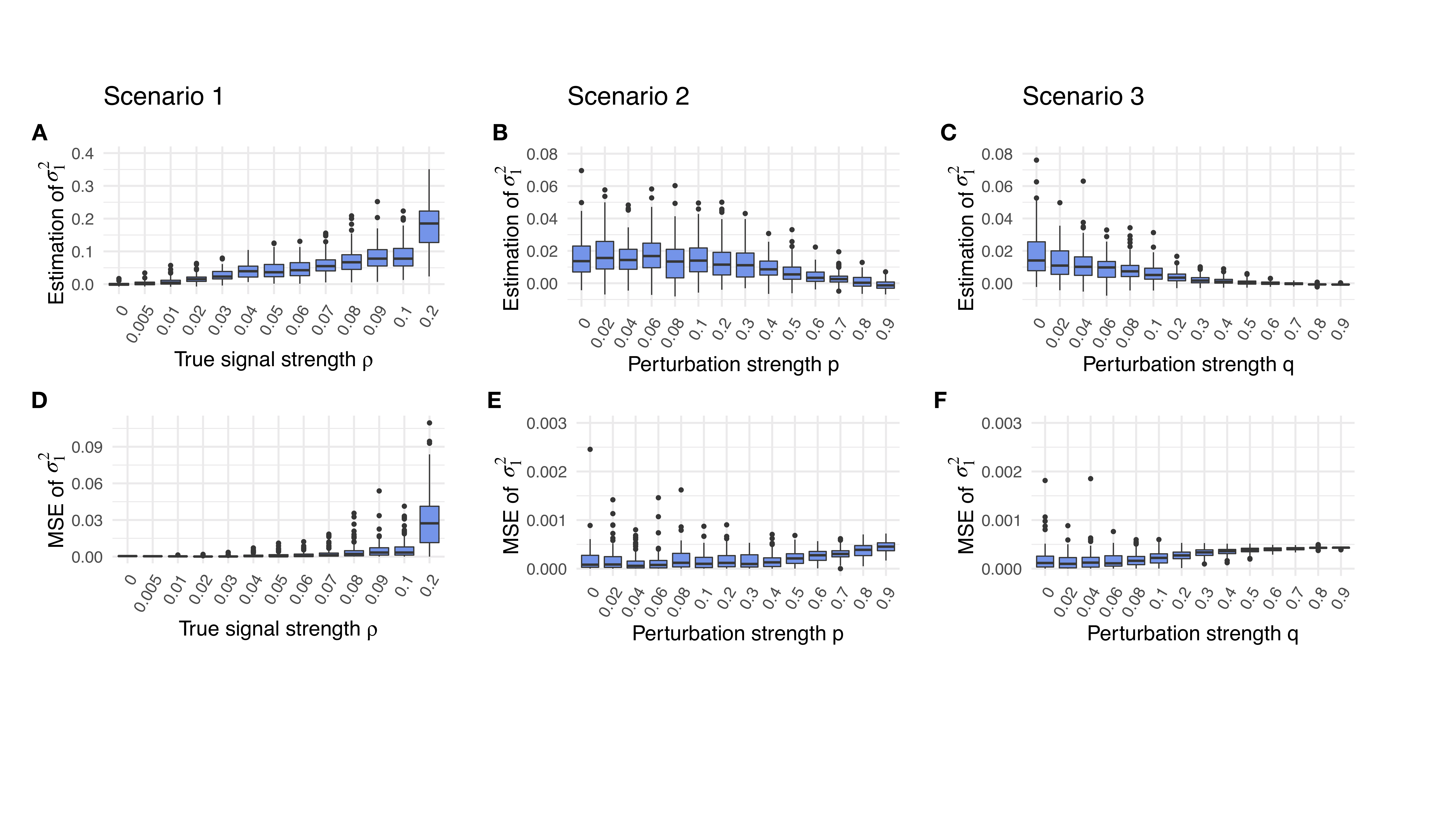

**S4 Fig. Parameter estimation of the parameter** $\boldsymbol{\sigma}_{\boldsymbol{1}}^{\boldsymbol{2}}$ **in simulations.** (A): Boxplot shows the $\sigma_{1}^{2}$ estimates across 100 simulation replicates for each signal strength parameter $\rho$ (x-axis) in the simulation scenario I. (B): Boxplot shows the $\sigma_{1}^{2}$ estimates across 100 simulation replicates for each parameter *p* (x-axis) in the simulation scenario II. Here, true $\sigma_{1}^{2}=0.02$. Increasing parameter *p* adds increasingly large noise to the tissue-specific adjacency matrices, but does not appear to strongly influence the estimation of $\sigma_{1}^{2}$. (C): Boxplot shows the $\sigma_{1}^{2}$ estimates across 100 simulation replicates for each parameter *q* (x-axis) in the simulation scenario III. Here, true $\sigma_{1}^{2}=0.02$. Increasing parameter *q* adds increasingly large noise to the tissue-specific adjacency matrices, but does not appear to strongly influence the estimation of $\sigma_{1}^{2}$. (D-F): mean squared error (MSE; y-axis) measures the accuracy of $\sigma_{1}^{2}$ estimates across the three simulation scenarios.

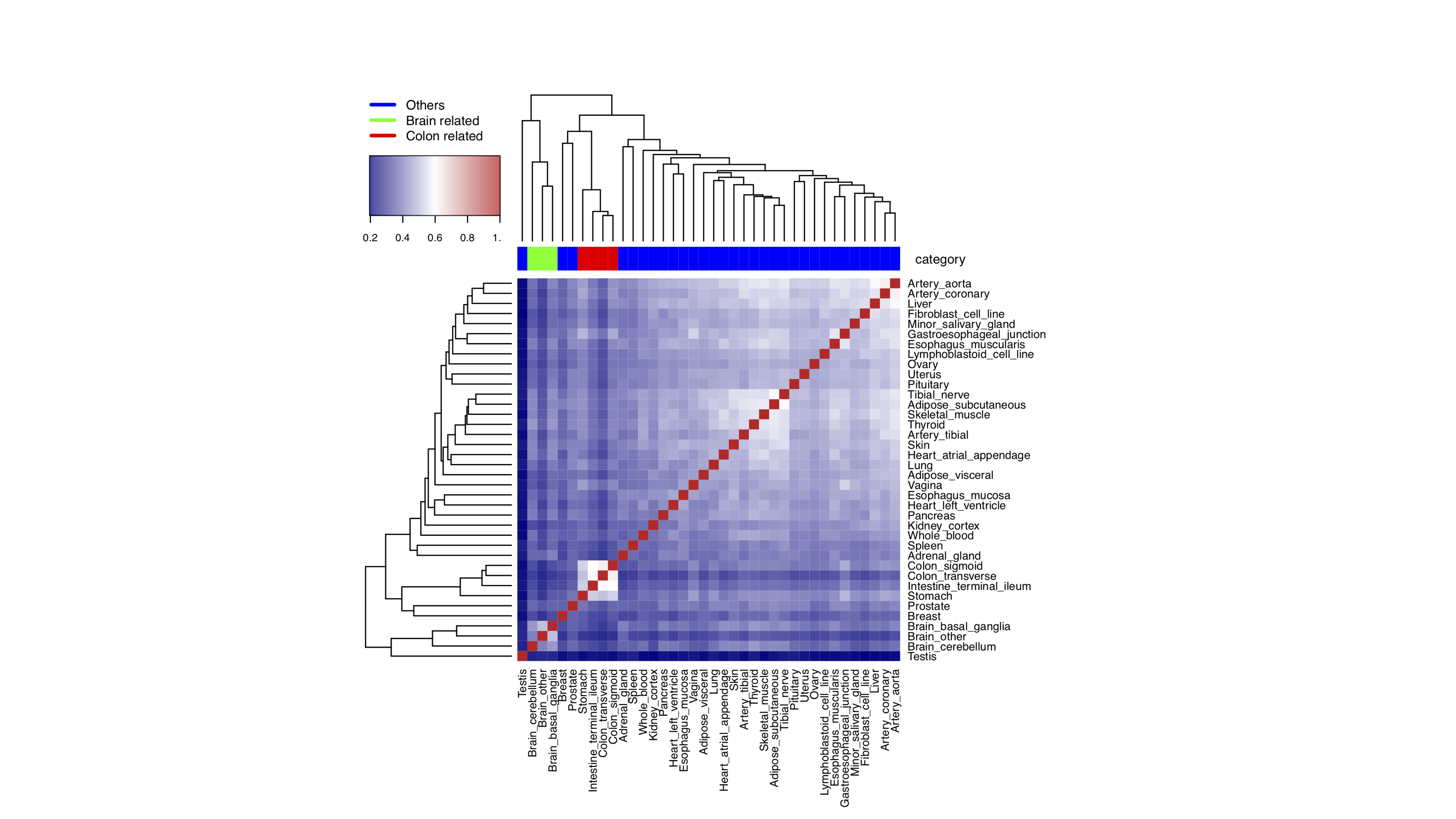

**S5 Fig. Adjacency matrices constructed from 38 tissues in the GTEx RNAseq data display tissue specific information.** Jaccard index is computed between pairs of matrices to measure the similarity among adjacency matrices across tissues. Adjacency matrices on similar tissues tend to cluster together based on hierarchical clustering. For example, the adjacency matrices for the three brain tissues (green), such as basal ganglia, cerebellum, and brain other, are all clustered together. Similarly, intestinal tissues (red), such as stomach, colon-transverse, intestine terminal ileum, and colon sigmoid, are all clustered together.

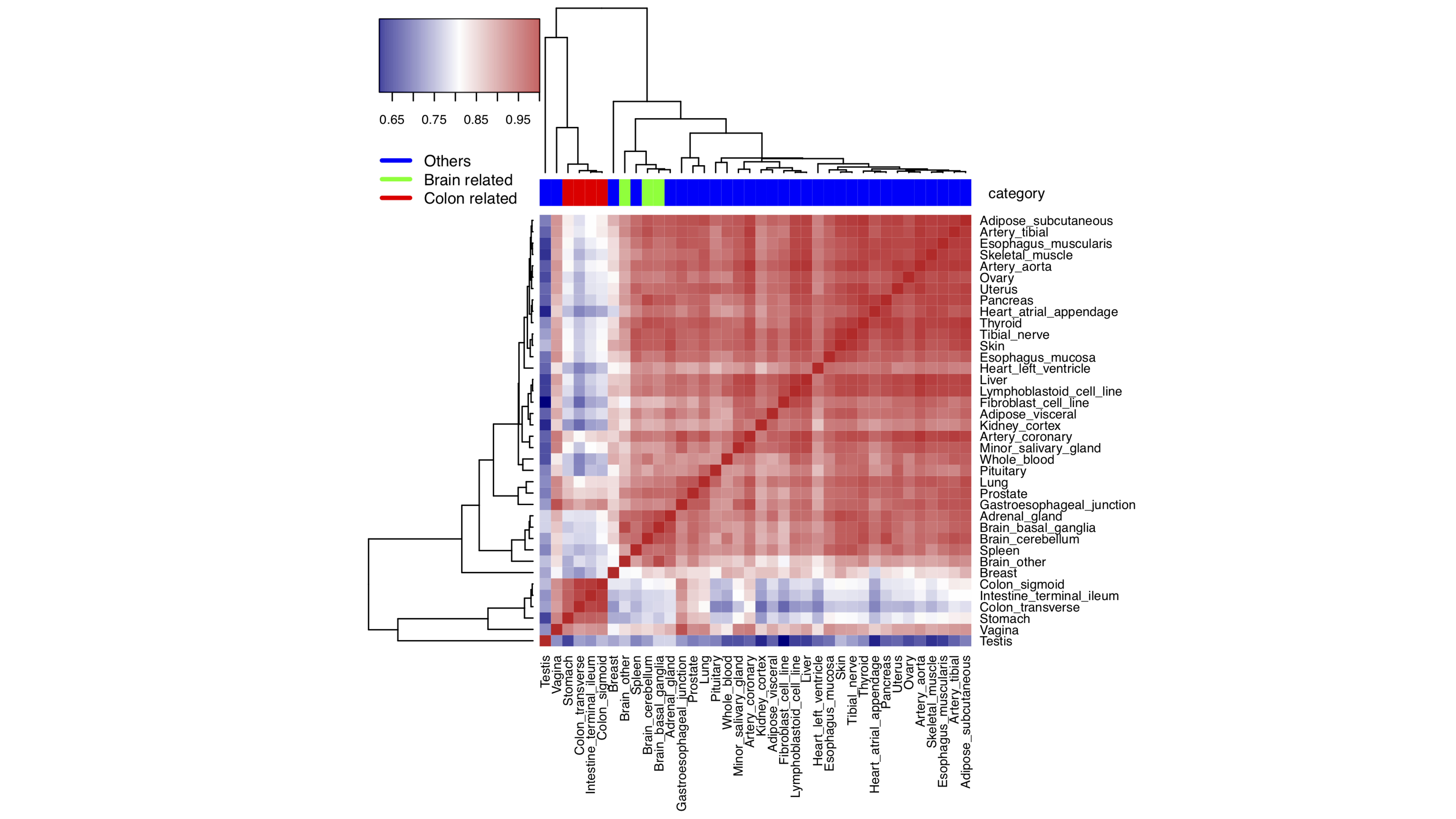

**S6 Fig. Gene connectivity in the adjacency matrices constructed from 38 tissues in the GTEx RNAseq data displays tissue specific information.** In each tissue, we calculated for each gene a node connectivity value, which measures the number of genes it is directly connected to a gene with a high node connectivity value is often referred to as a hub gene. Node connectivity values are similar between similar tissues as measured by Pearson’s correlation; thus similar tissues tend to cluster together based on Pearson’s correlation.

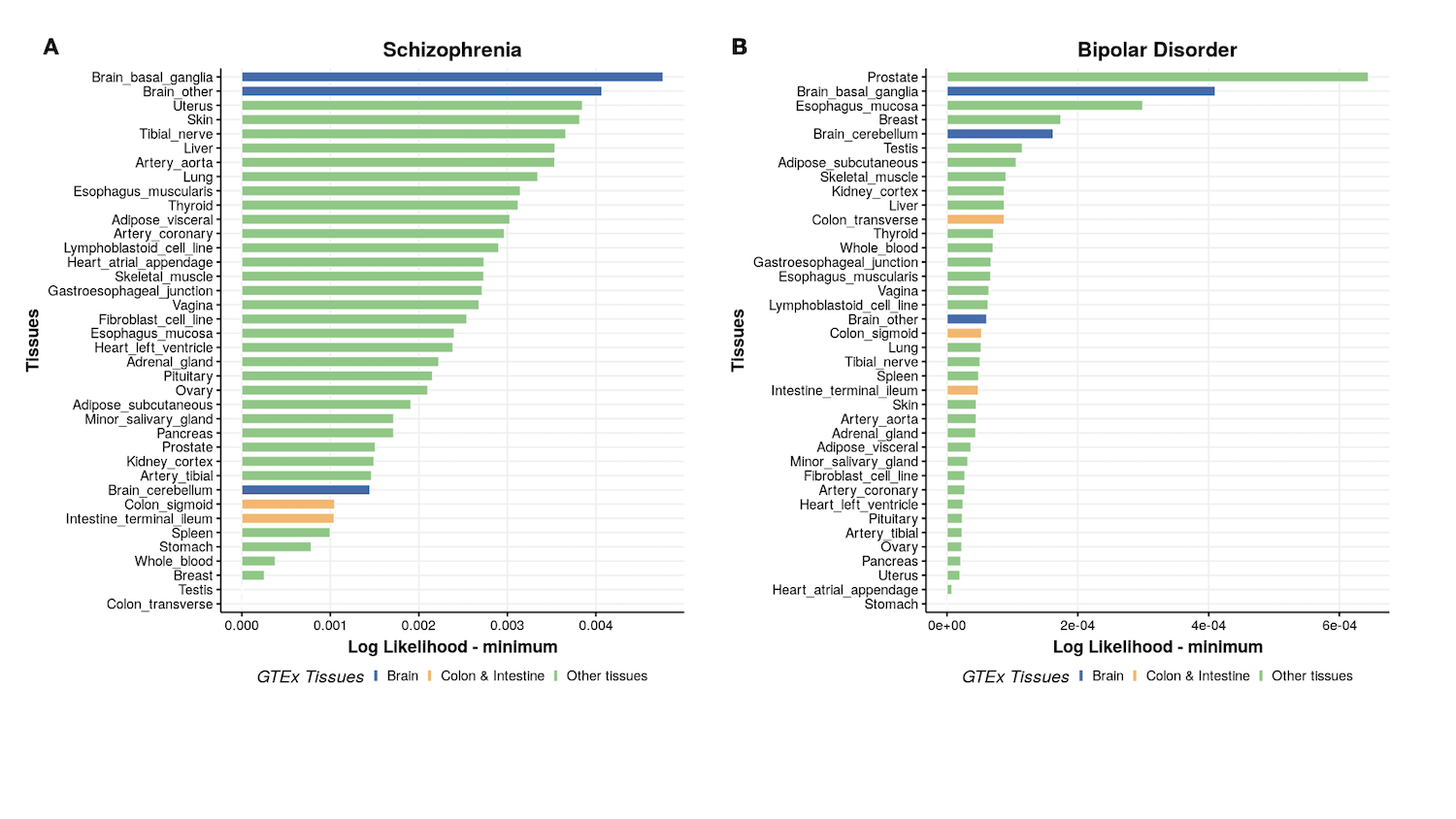

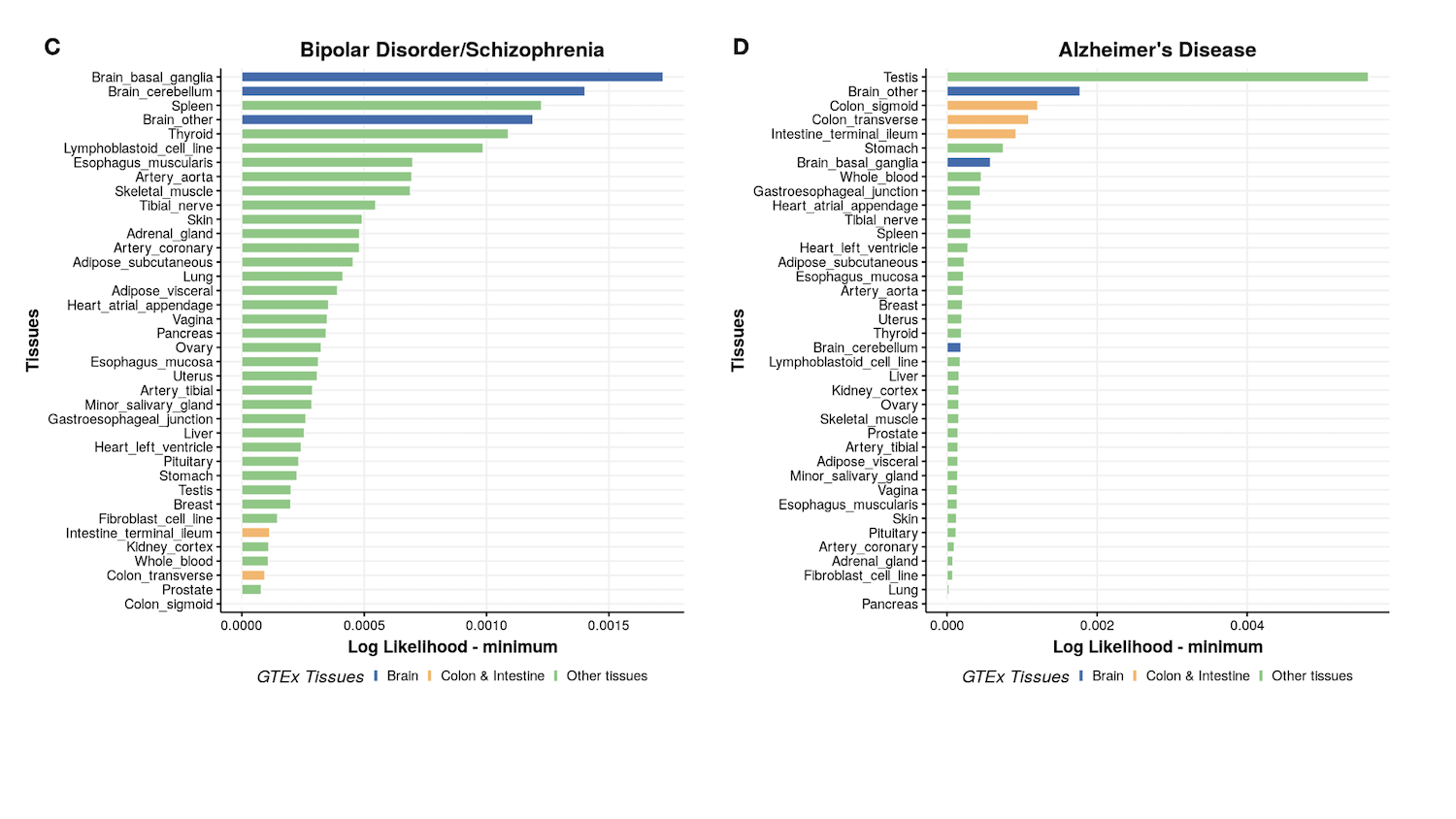

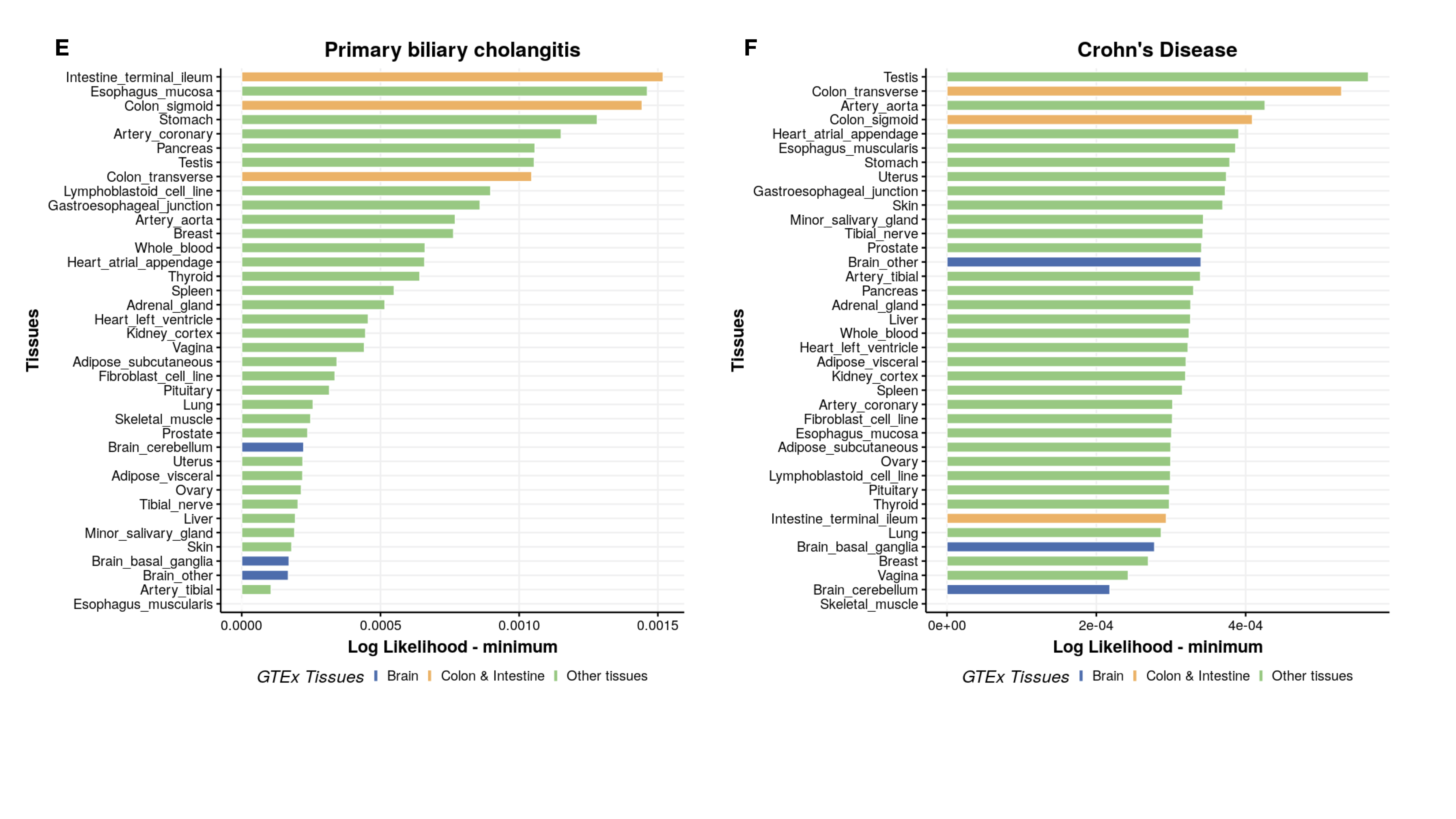

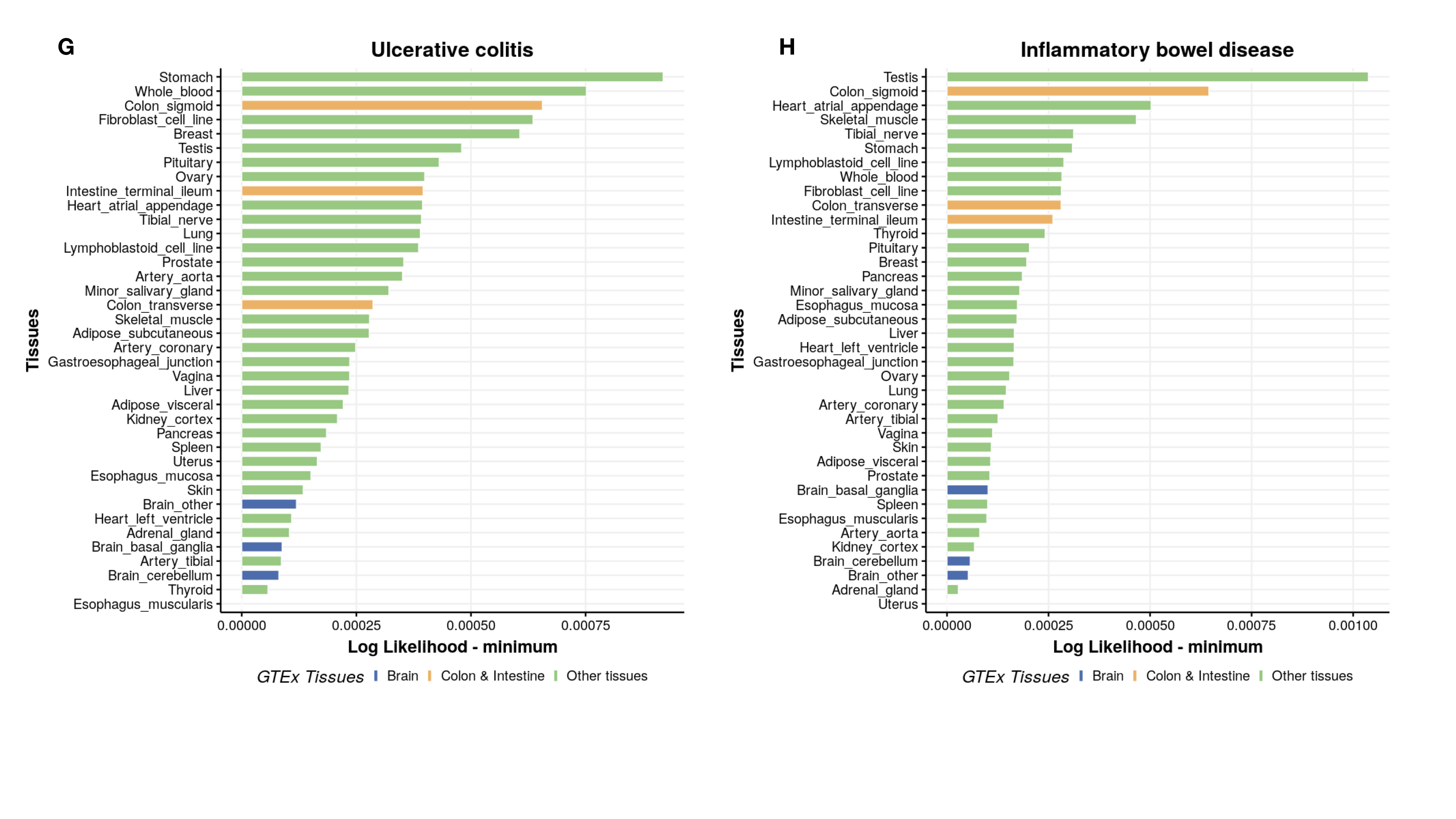

**S7 Fig. Rank of 38 tissues in the GTEx data in terms of their relevance to each of the eight GWAS traits obtained using CoCoNet.** For each GWAS trait, we calculated the composite likelihood for each tissue, subtracted the minimum likelihood across all tissues (x-axis), and ranked traits based on these values from top to bottom in each panel (y-axis). The brain tissues are colored in blue; the colon related tissues are colored in yellow; and the rest of the tissues are colored in green. Brain tissues tend to rank high for the four neurological diseases (top four panels) while colon related tissues tend to rank high for autoimmune diseases (bottom four panels).

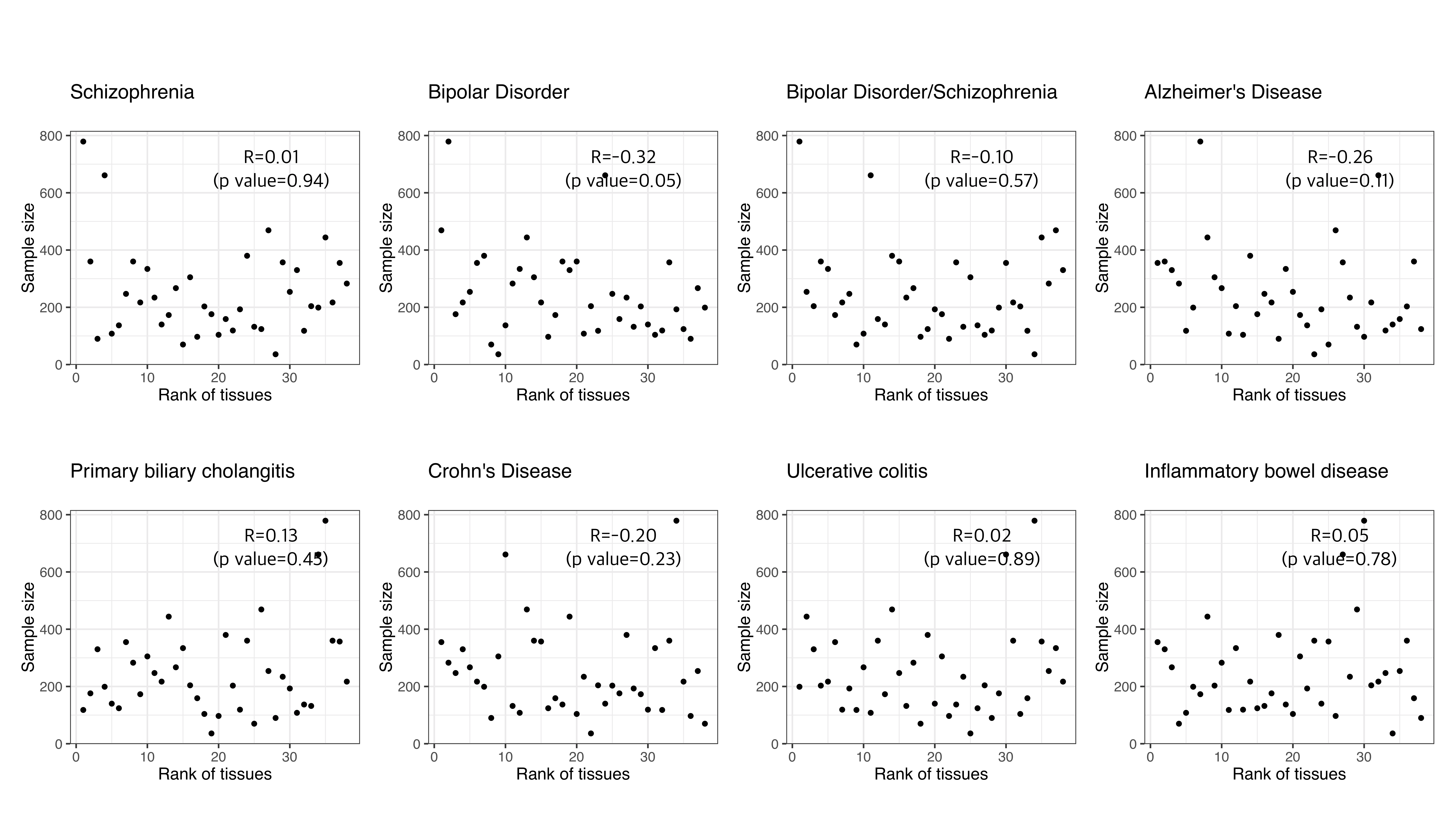

**S8 Fig. Tissue rank does not depend on tissue sample size for GWAS traits.** The sample size of each tissue (y-axis) is plotted against the rank of each tissue by CoCoNet (x-axis) across eight GWAS traits (eight panels). Spearman’s rank correlation (R) between the tissue rank and the sample size, together with the corresponding *p*-value, are also displayed on the panels. The tissue rank obtained by CoCoNet is not correlated with tissue sample size for all traits.

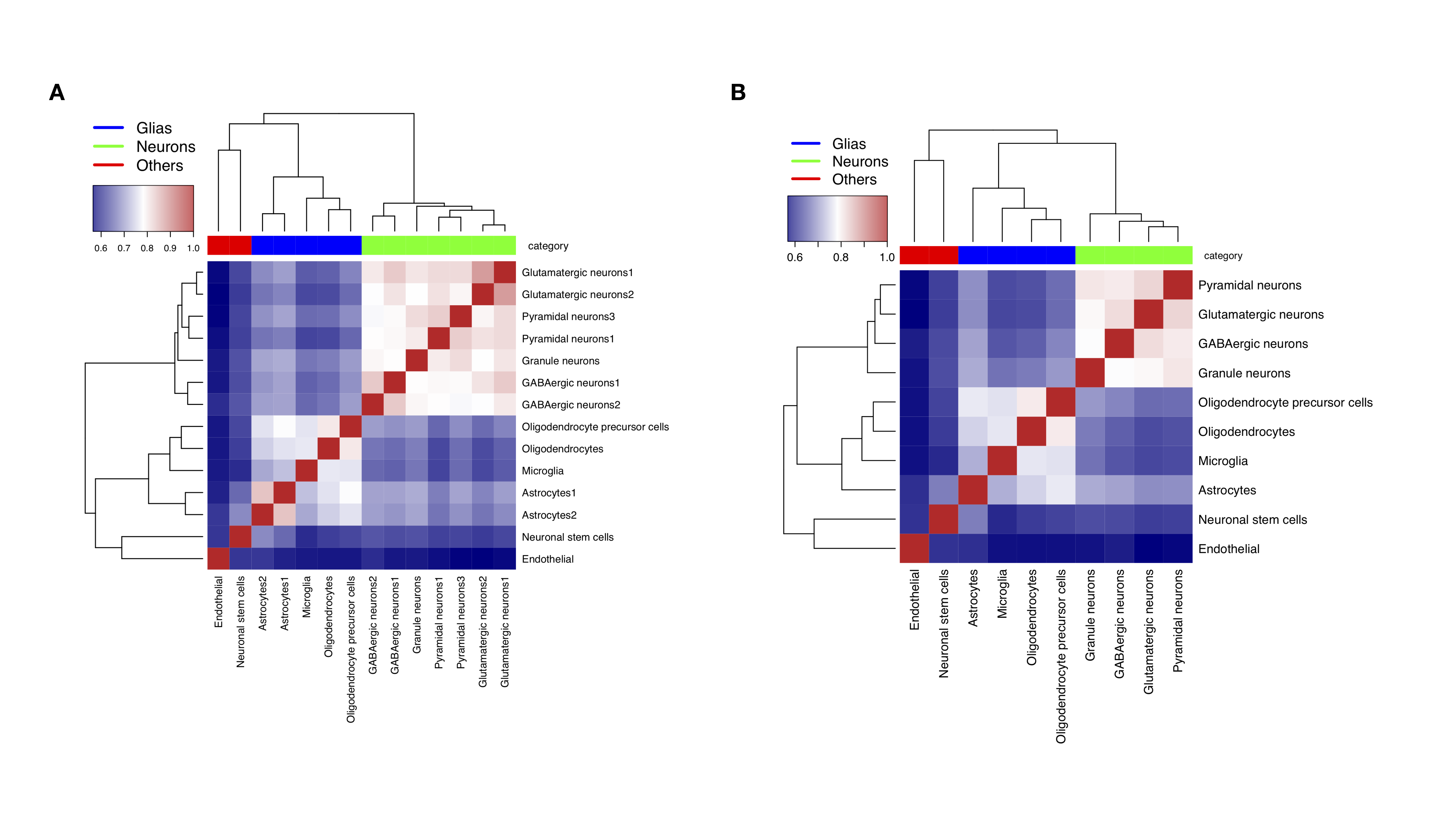

**S9 Fig. Heatmap showing Jaccard index** **among adjacency matrices constructed from multiple cell types in the single cell RNAseq data.** Jaccard index between adjacency matrices constructed either from cell types in two donors separately (**A**) or from cell types merged from two donors (**B**) is measured by Jaccard index. Adjacency matrices on similar tissues tend to cluster together based on hierarchical clustering. For example, the same cell type from different donors tend to cluster together (**A**) and different glia cell types tend to cluster together (**B**). The Jaccard index between two identical matrices is 1, as shown on the diagonal with red color in the heatmap.

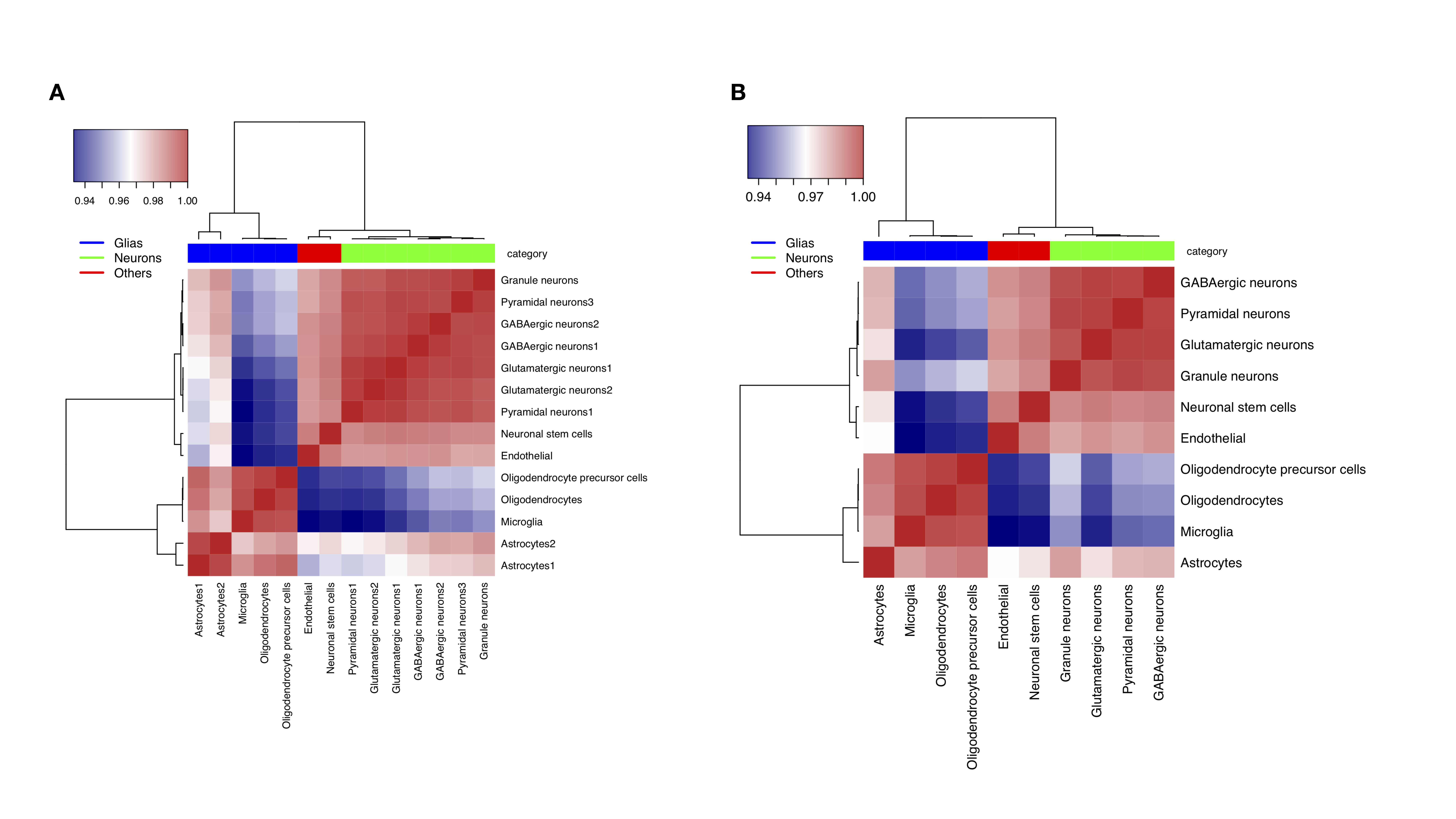

**S10 Fig. Gene connectivity in the adjacency matrices constructed from multiple cell types in the single cell RNAseq data.** We calculated for each gene a node connectivity value, which measures the number of genes it is directly connected to a gene with a high node connectivity value is often referred to as a hub gene. Similarity between cell types constructed either from two donors separately (**A**) or from cell types merged from two donors (**B**) is measured by Pearson’s correlation. The Pearson’s correlation between two identical matrices is 1, as shown on the diagonal with red color in the heatmap.

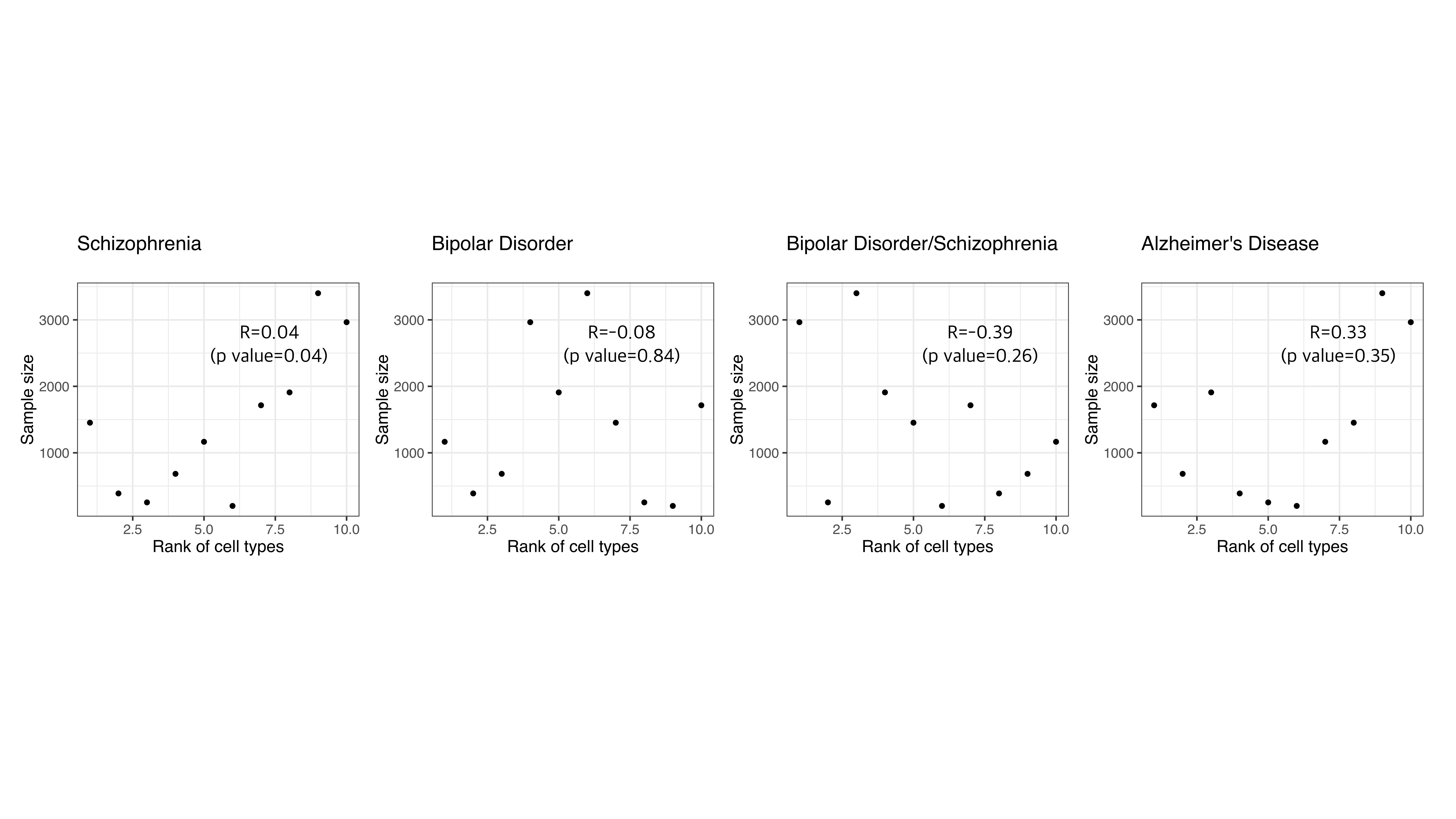

**S11 Fig. Cell type rank does not depend on the number of cells in each cell type for GWAS traits.** The number of cells in each tissue (y-axis) is plotted against the rank of each cell type by CoCoNet (x-axis) across eight GWAS traits (eight panels). Spearman’s rank correlation (R) between the cell type rank and the number of cells, together with the corresponding *p*-value, are also displayed on the panels. The cell type rank obtained by CoCoNet is not correlated with the number of cells in the cell type for all traits.

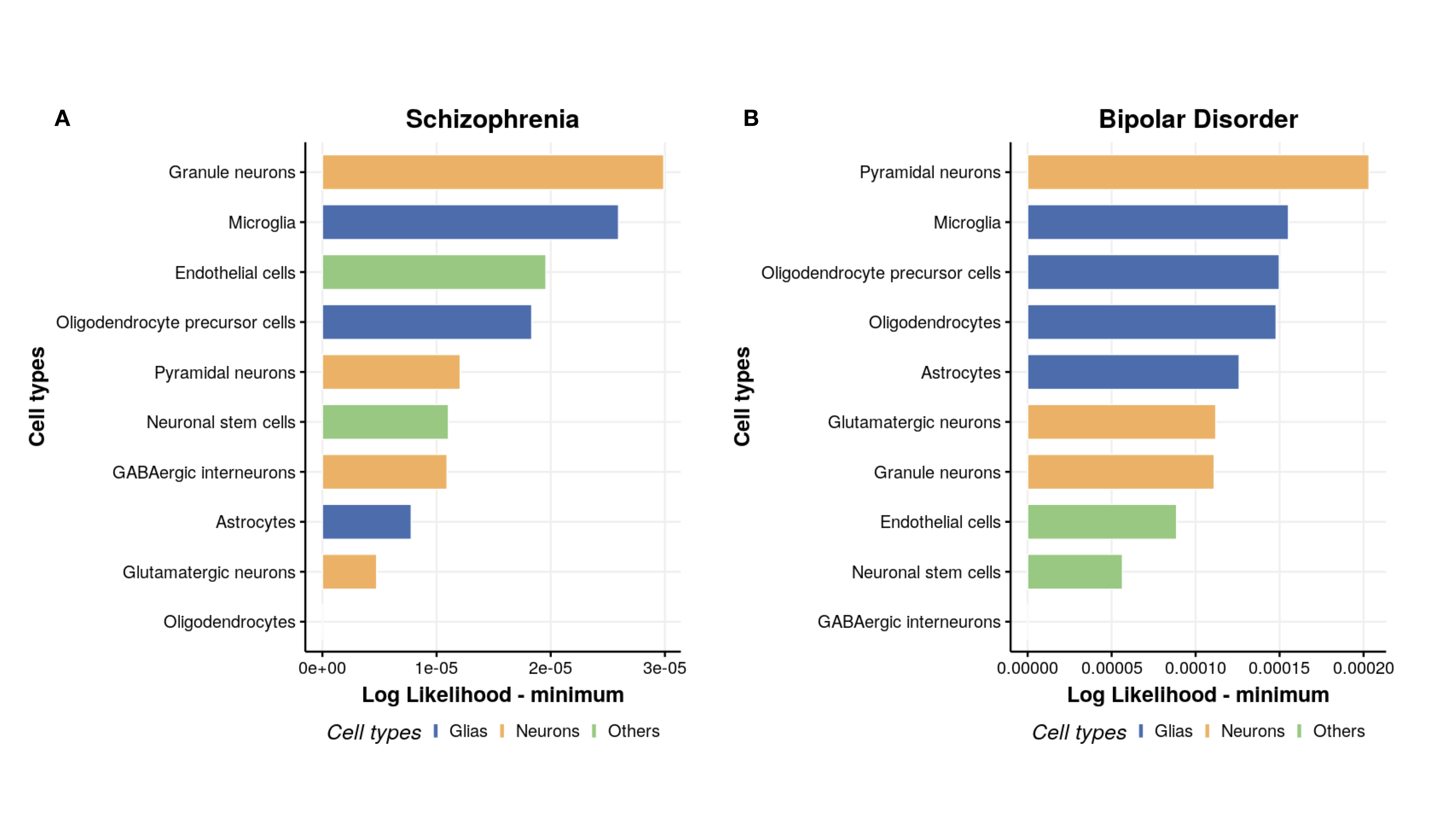

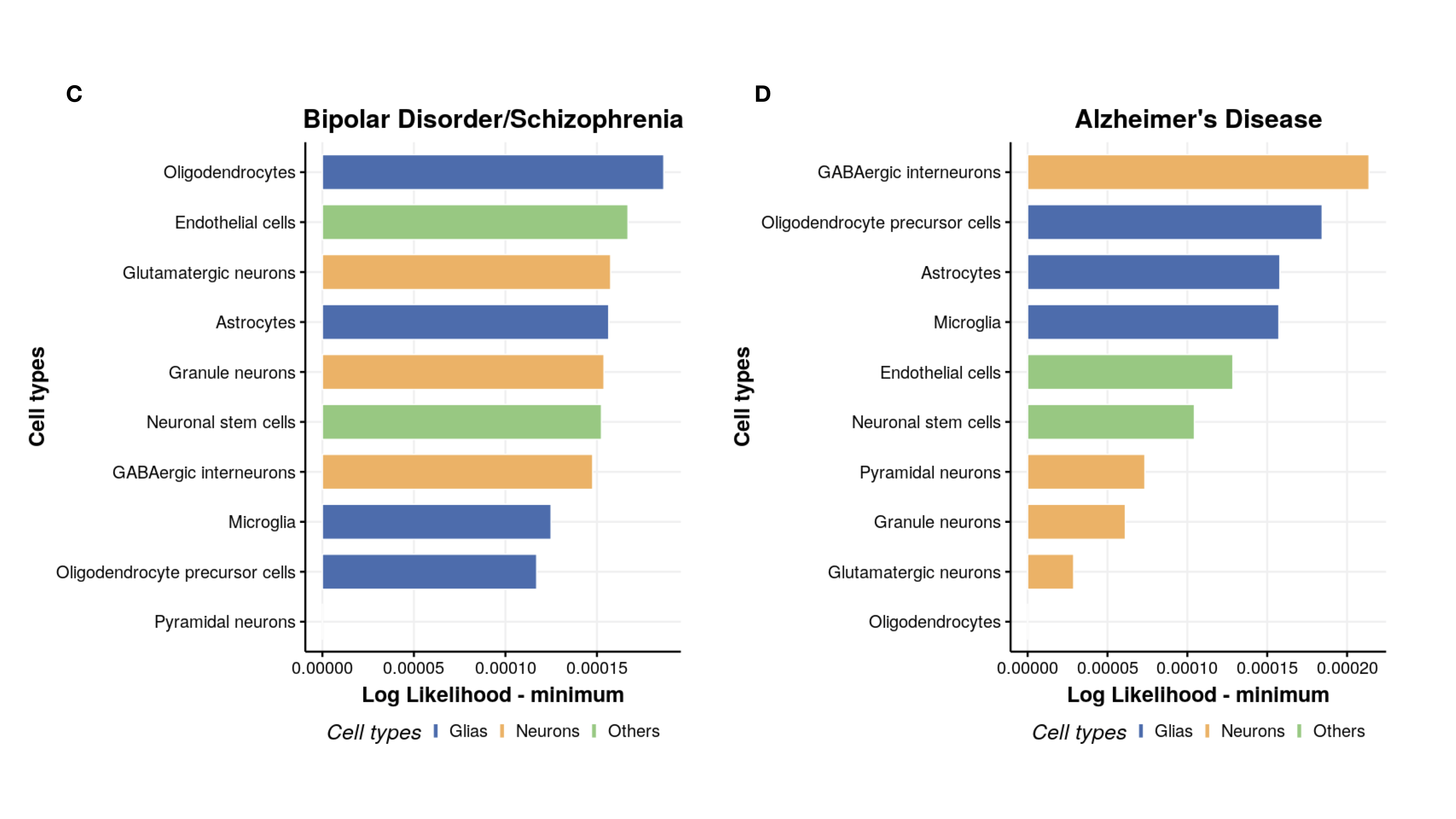

**S12 Fig. Rank of 10 cell types in terms of their relevance for each of the four neurological diseases obtained through CoCoNet.** For each of the four GWAS trait, we calculated the composite likelihood for each cell type, subtracted the minimum likelihood across all cell types (x-axis), and ranked cell types based on these values from top to bottom in each panel (y-axis).

**S1 Table. Computation time for CoCoNet in the two real data applications.**

Computing time is based on analysis of each trait-tissue pair using a single thread on a Xeon CPU E5-2620 v2 @ 2.10GHz processor.

| Datasets | Gene number | Time |
| --- | --- | --- |
| GTEx Tissues | 5359 | On average 7.2 min (min=5.2, max=11.6) |
| GTEx Cells | 8269 | On average 16 min (min=12.5,max=21.3) |

**S2 Table. Information for the summary statistics of eight GWAS traits.**

| No | Phenotype | Study Category | Abbreviation | Number of Samples | Reference | Web |
| --- | --- | --- | --- | --- | --- | --- |
| 1 | Schizophrenia | Neurological | SCZ | 70100 | (Ripke et al., 2014) | <http://www.med.unc.edu/>  pgc/downloads/ |
| 2 | Bipolar disorder | Neurological | BIP | 16731 | (Ruderfer et al., 2014) | <http://www.med.unc.edu/>  pgc/downloads/ |
| 3 | Bipolar disorder & Schizophrenia | Neurological | BIPSCZ | 39202 | (Ruderfer et al., 2014) | <http://www.med.unc.edu/>  pgc/downloads/ |
| 4 | Alzheimer's disease | Neurological | Alzheimer | 54162 | (Lambert et al., 2013) | <https://data.broadinstitute.org/>  alkesgroup/sumstats_formatted/ |
| 5 | Primary Biliary Cholangitis | Immune | PBC | 13239 | (Cordell et al., 2015) | <https://www.immunobase.org/>  downloads/protected_data/  GWAS_Data/ |
| 6 | Crohn's disease | Immune | CD | 20883 | (Jostins et al., 2012a) | <http://www.ibdgenetics.org/>  downloads.html |
| 7 | Ulcerative colitis | Immune | IBD | 34652 | (Jostins et al., 2012a) | <http://www.ibdgenetics.org/>  downloads.html |
| 8 | Inflammatory bowel disease | Immune | UC | 27432 | (Jostins et al., 2012a) | <http://www.ibdgenetics.org/>  downloads.html |

The table lists the phenotype name, phenotype category, abbreviation, number of individuals, reference, and downloaded websites for each of the eight GWAS traits.

**S3 Table. PubMed search keywords for eight GWAS traits.**

| No | Traits | keywords |
| --- | --- | --- |
| 1 | Schizophrenia | Schizophrenia[Title/Abstract] |
| 2 | Bipolar disorder | Bipolar disorder[Title/Abstract] |
| 3 | Bipolar disorder & Schizophrenia | (Bipolar disorder Schizophrenia[Title/Abstract] OR Bipolar disorder[Title/Abstract] OR Schizophrenia[Title/Abstract]) |
| 4 | Alzheimer's disease | (Alzheimer's disease[Title/Abstract] OR Alzheimer[Title/Abstract]) |
| 5 | Primary Biliary Cholangitis | (Primary Biliary Cholangitis[Title/Abstract] OR PBC[Title/Abstract]) |
| 6 | Crohn's disease | (Crohn's disease[Title/Abstract] OR Crohn[Title/Abstract]) |
| 7 | Ulcerative colitis | (Ulcerative colitis[Title/Abstract] OR Ulcerative[Title/Abstract]) |
| 8 | Inflammatory bowel disease | (Inflammatory bowel[Title/Abstract] OR IBD[Title/Abstract]) |

**S4 Table. PubMed search keywords for 38 tissues.**

| No. | Tissue name | Keywords |
| --- | --- | --- |
| 1 | adipose subcutaneous | (adipose subcutaneous[Title/Abstract] OR subcutaneous adipose[Title/Abstract]) |
| 2 | adipose visceral | (adipose visceral[Title/Abstract] OR visceral adipose[Title/Abstract]) |
| 3 | adrenal gland | adrenal gland[Title/Abstract] |
| 4 | artery aorta | artery aorta[Title/Abstract] |
| 5 | artery coronary | artery coronary[Title/Abstract] |
| 6 | artery tibial | artery tibial[Title/Abstract] |
| 7 | Brain other | (substantia nigra[Title/Abstract] OR hypothalamus[Title/Abstract] OR hippocampus[Title/Abstract] OR frontal lobe[Title/Abstract] OR cerebral cortex[Title/Abstract] OR amygdala[Title/Abstract]) |
| 8 | Brain cerebellum | cerebellum[Title/Abstract] |
| 9 | Brain basal ganglia | (nucleus accumbens[Title/Abstract] OR caudate putamen[Title/Abstract] OR caudate nucleus[Title/Abstract]) |
| 10 | breast | breast[Title/Abstract] |
| 11 | lymphoblastoid cell line | lymphoblastoid cell line[Title/Abstract] |
| 12 | fibroblast cell line | fibroblast cell line[Title/Abstract] |
| 13 | colon sigmoid | (colon sigmoid[Title/Abstract] OR large intestine[Title/Abstract]) |
| 14 | colon transverse | (colon transverse[Title/Abstract] OR large intestine[Title/Abstract]) |
| 15 | gastroesophageal junction | gastroesophageal junction[Title/Abstract] |
| 16 | esophagus mucosa | esophagus mucosa[Title/Abstract] |
| 17 | esophagus muscularis | esophagus muscularis[Title/Abstract] |
| 18 | heart atrial appendage | heart atrial appendage[Title/Abstract] |
| 19 | heart left ventricle | heart left ventricle[Title/Abstract] |
| 20 | kidney cortex | kidney cortex[Title/Abstract] |
| 21 | liver | liver[Title/Abstract] |
| 22 | lung | lung[Title/Abstract] |
| 23 | minor salivary gland | minor salivary gland[Title/Abstract] |
| 24 | skeletal muscle | skeletal muscle[Title/Abstract] |
| 25 | tibial nerve | tibial nerve[Title/Abstract] |
| 26 | ovary | ovary[Title/Abstract] |
| 27 | pancreas | pancreas[Title/Abstract] |
| 28 | pituitary | pituitary[Title/Abstract] |
| 29 | prostate | prostate[Title/Abstract] |
| 30 | skin | skin[Title/Abstract] |
| 31 | intestine terminal ileum | (intestine terminal ileum[Title/Abstract] OR terminal ileum[Title/Abstract]) |
| 32 | spleen | spleen[Title/Abstract] |
| 33 | stomach | stomach[Title/Abstract] |
| 34 | testis | testis[Title/Abstract] |
| 35 | thyroid | thyroid[Title/Abstract] |
| 36 | uterus | uterus[Title/Abstract] |
| 37 | vagina | vagina[Title/Abstract] |
| 38 | whole blood | whole blood[Title/Abstract] |

**S5 Table. PubMed search keywords for 10 cell types.**

| No | Cell type name | keywords |
| --- | --- | --- |
| 1 | ASC | astrocytes[Title/Abstract] |
| 2 | END | endothelial[Title/Abstract] |
| 3 | GABA | GABAergic[Title/Abstract] |
| 4 | MG | microglia[Title/Abstract] |
| 5 | NSC | neuronal stem cells[Title/Abstract] |
| 6 | ODC | oligodendrocytes[Title/Abstract] |
| 7 | OPC | oligodendrocyte precursor cells[Title/Abstract] |
| 8 | exCA | pyramidal neurons[Title/Abstract] |
| 9 | exDG | granule neurons[Title/Abstract] |
| 10 | exPFC | glutamatergic neurons[Title/Abstract] |
